# Periodontopathogens interfere with the human renin-angiotensin system through surface-attached proteases

**DOI:** 10.1101/2024.06.20.599981

**Authors:** Irena Waligórska, Krzysztof Żak, Joanna Budziaszek, Ewa Bielecka, Tomasz Kantyka, Joanna Kozieł, Ida B. Thøgersen, Jan J. Enghild, Przemysław Grudnik, Jan Potempa, Mirosław Książek

## Abstract

The renin-angiotensin system (RAS) executes its functions through biologically active peptides, angiotensins (Ang). Angiotensinogen-derived precursor, Ang I is cleaved by angiotensin-converting enzyme (ACE) into proinflammatory Ang II, which increases blood pressure. In the alternate pathway performed by neprilysin and ACE II, Ang 1-7 is produced from Ang I with activities opposite to Ang II. Here, we show that *Porphyromonas gingivalis* (*Pg*) and *Tannerella forsythia* (*Tf*), endogenous oral pathogens, direct RAS into generation of Ang 1-7 through endopeptidases O, PgPepO and TfPepO, respectively. PepOs are thermophilic metalloproteases inhibited by cation chelators, but not by specific ACE and neprilysin inhibitors. PgPepO and TfPepO prefer large hydrophobic amino acids at the carbonyl side of scissile peptide bonds (P1’ position), and TfPepO, contrary to all known homologous proteases, hydrolyzes substrates away from both terminuses. Solved crystal structures show that exceptionally wide entrance to the catalytic cleft explains unique properties of TfPepO. Furthermore, the different nature of subsites S1’ and S2’ in the substrate binding site explains refractory of PepOs to inhibitors of human homologous proteases. Multiple immunoassays clearly show that PepOs are attached to the bacteria cell surface and are released in outer membrane vesicles. Moreover, PepO is responsible for Ang I hydrolysis by *Pg* and *Tf*. Finally, PepO deletion reduced only the virulence of *Tf* in the *Galleria mellonella* model. Thus, our data show that *Pg* and *Tf* interfere with RAS through a PepO protease.

## Introduction

The human body has many systems responsible for its proper functioning. One is the hormonal-enzymatic renin-angiotensin system (RAS), that regulates blood pressure and water and electrolyte balance. The two main components of the RAS system are a peptide hormone, angiotensin II (Ang II), and renin, an aspartic protease produced by kidney granular cells and hydrolyzing angiotensinogen circulating in the blood to angiotensin I (Ang I) (Crisan & Carr, 2000). Ang I is converted to other angiotensins during further processing by two pathways. The first involves the removal of two amino acids from the C terminus of Ang I by angiotensin-converting enzyme (ACE), resulting in Ang II formation. Through interaction with the AT_1_ receptor (AT_1_R), Ang II is a strongly pro-inflammatory and vasoconstrictive factor and also increases oxidative stress, hypertrophy and cell proliferation (Weber & Brilla, 1991; Campbell et al., 1995; Welch 2008). The second pathway, executed by ACE II and a neural endopeptidase, neprilysin, leads to the formation of angiotensin 1-7 (Ang 1-7) directly from Ang I or through the intermediate angiotensin 1-9 (Ang 1-9) (Rice et al., 2004; Becari et al., 2011; Ferreira et al., 2012). In contrast to Ang II, Ang 1-7 has anti-inflammatory effects, reduces oxidative stress and decreases blood pressure through interaction with Mas receptor (MasR) (Ferreira *et al.,* 2012; Mordwinkin *et al.,* 2012; Patel et al., 2016). Dysregulation of RAS may lead to cardiovascular diseases such as hypertension (Howell & Camerson, 2016) and other systemic conditions like atherosclerosis (Thomas et al., 2010) and diabetes (Zhang *et al*., 2015). In addition to its systemic role, components of RAS are also locally expressed in many tissues, such as periodontal tissues (Saravi et al., 2020), where the expression on the mRNA level of all critical elements of this system, presence of angiotensins (Santos et al., 2015) and interaction of Ang II with AT_1_R was shown (Nakamura et al., 2011).

Periodontitis, or periodontal disease, damages the tissues surrounding and supporting teeth and affects up to 20% of the world’s adult population in its severe forms (Nazir et al., 2020). If left untreated, the disease not only leads to tooth loss but also contributes to the progression and/or development of systemic diseases such as diabetes, rheumatoid arthritis, cardiovascular and neurodegenerative diseases and cancer (Paturel et al., 2004; Behle & Papapanou, 2006; Wegner et al., 2010; Sfyroeras et al., 2012; Chen *et al.,* 2017; de Miguel-Infante et al., 2018). Moreover, the disease causes significant health and economic burdens: in 2018 alone, the costs incurred due to periodontitis were estimated at USD 154.06 billion in the USA and EUR 158.64 billion in the European Union (Botelho et al., 2022). Periodontitis has a multifactorial, complex etiology, but the critical event leading to the development of the disease is the disruption of the homeostasis of the oral microflora in subgingival plaque (Saini et al., 2011). Plaque becomes pathogenic when the composition of the microbial community shifts towards a greater proportion of gram-negative anaerobic bacteria. The factor that causes such a change is the colonization of the plaque by the so-called “keystone pathogens” that can modulate the host response and, as a result, increase the virulence of the entire community. One such agent is *Porphyromonas gingivalis*, which is almost always found together with *Treponema denticola* and *Tannerella forsythia*. (Grenier et al., 1987; Hajishengallis & Lamont, 2012; Darveau et al., 2012). The presence of these three gram-negative anaerobic pathogens, forming a “red complex” of bacteria, is very strongly correlated with both; the occurrence of the disease and the severity of its symptoms (van Winkelhoff et al., 2002; Holt SC & Ebersole, 2005; Haffajee et al., 2008).

A common feature of “red complex” bacteria is the secretion of an array of extracellular proteases. *P. gingivalis* gingipains R (RgpA, RgpB) and K (Kgp), responsible for 85% of the extracellular proteolytic activity of this bacterium, are involved in nutrients acquisition, avoiding the host’s innate immunity and immunomodulation (Imamura 2003; Potempa & Pike, 2005; Guo et al., 2010; Bengtsson et al., 2015). *T. forsythia* secretes, among others, six KLIKK proteases (three serine proteases: mirolase, miropsin-1 and miropsin-2, and three metalloproteases: mirolysin, karilysin and forsilysin) sharing a unique multi-domain structure (Ksiazek et al., 2015). Karilysin and mirolysin are responsible for complement evasion and inactivation of LL-37, a crucial antimicrobial peptide in the human oral cavity (Karim et al., 2010; Koziel et al., 2010; Koneru et al., 2017). *P. gingivalis* and *T. forsythia* also secrete endopeptidases O (PepOs): PgPepO and TfPepO, respectively (Ansai et al., 2003; Friedrich et al., 2015). PgPepO is involved in the invasion of host cells (Ansai et al., 2003; Park et al., 2004) and cleaves big endothelin-1 and Ang I (Carson et al., 2002; Awano et al., 1999). However, knowledge about PgPepO is still minimal and there is no available data about TfPepO protease.

Herein, we report that periodontopathogens *P. gingivalis* and *T. forsythia* interfere with RAS through a single protease, PgPepO and TfPepO, respectively. We present a thorough biochemical characterization of the proteases, revealing their substrate specificity. Furthermore, we used X-ray crystallography to solve the structures of PepOs, which explains the unique properties of TfPepO. Moreover, we employ different methods to show that both PepOs are secretory lipoproteins attached to the cell surface. Finally, the *in vivo* model uncovers the different roles of PepOs in pathogens’ evasion of innate immunity.

## Results

### PgPepO and TfPepO are putative lipoproteins from the M13 family of metalloproteases

PepO proteases found in the genome of *P. gingivalis* (PgPepO) and *T. forsythia* (TfPepO) belong to the MEROPS M13 family of proteases, the progenitor of which is human neprilysin (Rawlings et al., 2018). Sequence alignment of PgPepO and TfPepO with six other members of the M13 family, three human and three bacterial enzymes showed that PgPepO and TfPepO contain a zinc-binding consensus sequence (ZBCS) consisting of two histidine residues and a glutamic acid residue located 25 amino acids residues towards the C-terminus from the main motif (HEXXH…E), in which the first Glu residue has a catalytic function in proteolysis (SI appendix, Fig. S1). In addition, human M13 proteases have disulfide bridges: six (hNEP) or five (human endothelin-converting enzyme 1, hECE1, and 2, hECE2). However, bacterial PepO probably does not have disulfide bridges, which was confirmed by solved crystal structures (PepO from the bacterium *Lactobacillus rhamnosis*, PDB accession no.: 4IUW) and the absence of conserved cysteines forming disulfide bridges in human representatives of the M13 family of proteases in prokaryotic PepO. PgPepO and TfPepO have a signal peptide characteristic of secretory lipoproteins with a length of 22 and 16 amino acid residues, respectively (Juncker et al., 2003). We obtained these two PepO from periodontopathogens as a recombinant proteins using the *Escherichia coli* expression system. The monomeric tag-free proteins were obtained by affinity chromatography on glutathione-Sepharose, with simultaneous removal of the GST tag using PreScission protease, and further by size exclusion chromatography (SI appendix, Fig. S2, S3).

### Biochemical characterization

The fluorescent substrate Mca-RPPGFSAFK(Dnp)-OH, whose amino acid sequence is based on the sequence of bradykinin (Johnson & Ahn, 2000) was used to determine the activity of human M13 metalloproteases (NEP, ECE1 and ECE2). We therefore used the same substrate to verify PgPepO and TfPepO proteolytic activity. Both proteases hydrolyzed Mca-RPPGFSAFK(Dnp)-OH, with K_m_ equal to 3 and 3.1 μM, respectively (Fig. 1A), while the catalytically inactive mutants, with Glu in the catalytic center replaced by Ala (PgPepO^E527A^ and TfPepO^E515A^), showed no activity towards this substrate (SI appendix, Figure S2G, S3F). Thus, this substrate was used for further biochemical characterization of PepOs.

**Figure 1.**
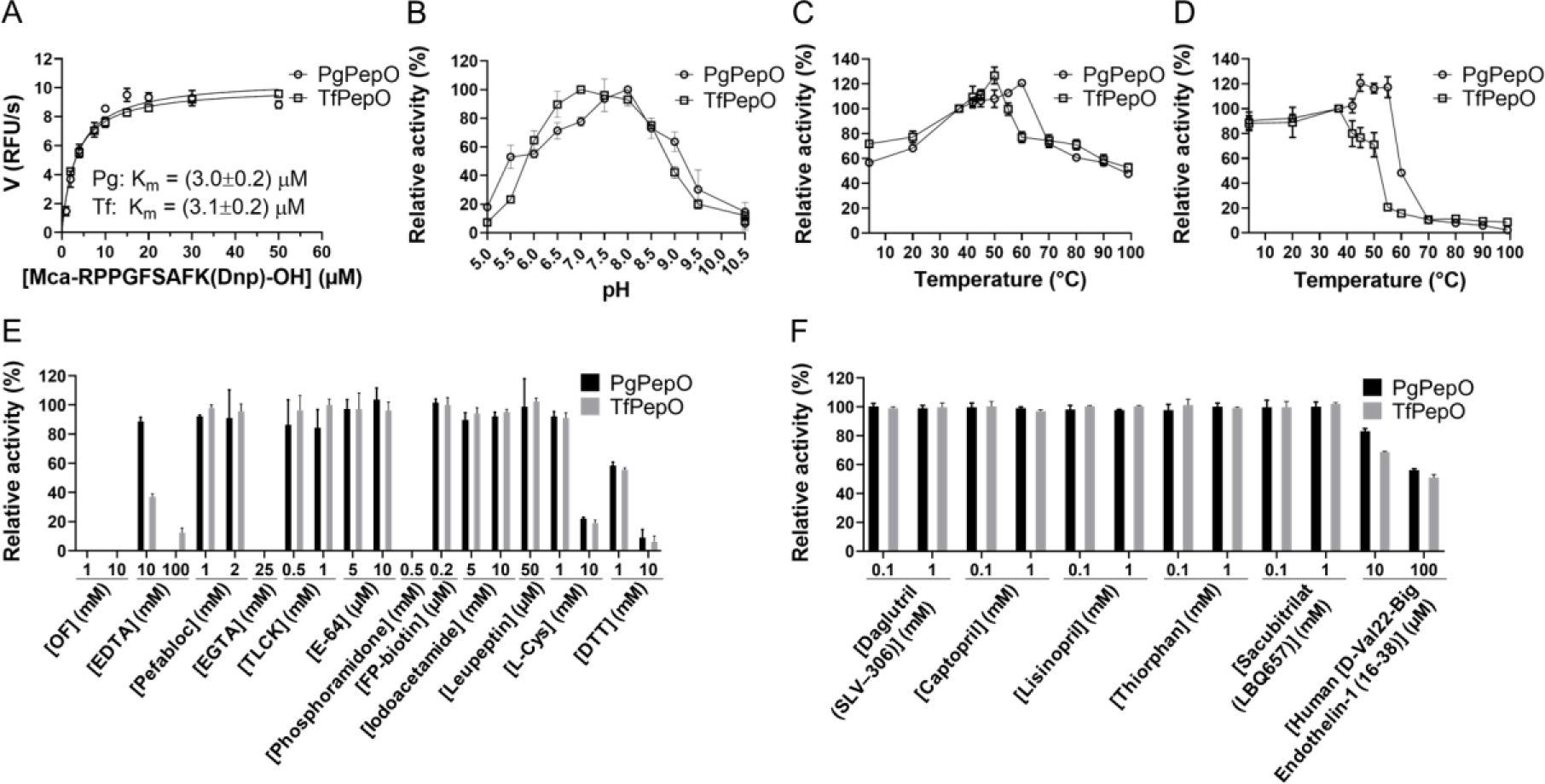
Biochemical characterization of PgPepO and TfPepO. A) Velocity of reaction (V) was determined for different concentrations of the substrate Mca-RPPGFSAFK(Dnp)-OH, and the Michaelis-Menten constant (K_m_) was calculated employing GraphPad Prism macro. (B, C) Effect of pH (B) and temperature (C) on PepOs activity. Enzymatic activity was measured at different pH (B) and temperatures (C). The pH with the highest V (B) and V at 37°C (C) was arbitrarily taken as 100%. (D) Thermal stability of PgPepO and TfPepO. PepOs were incubated at indicated temperatures for 30 min, and enzymatic activity was measured. Activity at 37°C was set at 100%. (E, F) The effect of inhibitors and reducing agents on the activity of PgPepO and TfPepO. PepOs were incubated with the indicated compound, and activity was measured. PepO activity determined at 37°C in 100 mM Tris, 150 mM NaCl, 0.02% NaN_3_, 0.05% Pluronic F-127 (pH 7.5) was arbitrarily taken as 100%. For all panels, data are means±SD (N=3).

First, we determined the pH and temperature optimum. The PepOs were active in a wide pH range (6-9), and they achieved the highest activity at pH 8 (PgPepO) and 7 (TfPepO) (Fig. 1B). PepOs were active throughout the entire temperature range tested (4-100°C). They achieved optimal activity at 60°C (PgPepO) and 50°C (TfPepO), and thus, they are thermophilic enzymes (Sterpone & Melchionna, 2012). However, both enzymes incubated alone for 30 min at 70°C lost virtually all activity (Fig. 1C, D). This discrepancy can be explained by the fact that the substrate bound to the PepOs increases their thermostability.

Second, we assessed the effect of various class-specific inhibitors and reducing agents on the PepOs’ activity. Inhibitors of metalloproteases inhibited PgPepO and TfPepO, including dipositive cations chelators (OF, EDTA, EGTA) and phosphoramidon, a competitive inhibitor of many metalloproteases (e.g. hNEP, hECE1 and thermolysin) (Rawlings et al., 2018). Inhibitors of serine (TLCK, Pefabloc, leupeptin, FP-biotin) and cysteine proteases (E-64, leupeptin, TLCK, iodoacetamide) had no effect on the activity of PgPepO and TfPepO (Fig. 1E). These results confirm the conclusions from bioinformatic analyzes that PgPepO and TfPepO are zinc-dependent metalloproteases. L-cysteine and DTT reduced the activity of both proteases even by 80% and 90%, respectively (Fig. 1E). This inhibition was likely caused by the binding of the free thiol group to the fourth coordination site of the catalytic zinc, which under normal conditions is occupied by a water molecule involved in proteolysis mechanism. Free thiol groups are used not only to develop metalloprotease inhibitors but also in the so-called “cysteine switch”, one of the zymogenicity mechanisms of metalloproteases (Müller et al., 1997).

Third, many metalloproteases bind an additional zinc ion termed the structural zinc, or other dipositive ions such as magnesium or calcium (Dudev & Lim, 2003). Thus, the influence of these three dipositive cations on the activity of PgPepO and TfPepO was checked. Mg^2+^ and Ca^2+^ cations (10 nM - 10 mM) had virtually no effect on the activity of PepOs. In the case of PgPepO, an increase in zinc concentration resulted in a gradual increase in enzyme activity by 20% until it reached a maximum at a concentration of 10 μM. However, at concentrations up to 10 μM, zinc ions had virtually no effect on the activity of TfPepO. A further increase in the concentration of zinc ions led to a gradual loss of activity of both PepOs and a complete loss of activity at a concentration of 10 mM (SI appendix, Figure S4).

Finally, due to their important physiological function, human enzymes from the M13 family: neprilysin, ECEs and RAS-related ACE, are important targets for the development of inhibitors. These compounds are often used in the treatment of hypertension. Therefore, the effect of the following compounds: Daglutril (SLV-306), Tiorfan and Sacubitrilat (LBQ657) (neprilysin inhibitors), Daglutril (SLV-306), Human [D-Val22]-Big Endothelin-1 (16-38) (ECE inhibitors) and Captopril, Lisinopril (ACE inhibitors) on PgPepO and TfPepO activity was checked (Fig. 1F). The inhibition was observed only by Human [D-Val22]-Big Endothelin-1 (∼50% at 100 μM inhibitor concentration).

### Substrate specificity

To determine substrate specificity, we checked whether PgPepO and TfPepO hydrolyze various proteins and peptides that are i) found in human fluids and thus can be digested at the site of infection, ii) degraded by the characterized metalloproteases from the M13 family or iii) widely used to determine the substrate specificity of proteolytic enzymes. The PepOs hydrolyzed three peptide substrates (human big endothelin-1, insulin B-chain, Ang 1-14) but not the antibacterial peptide LL-37, and only one proteinaceous substrate, α-casein. Other checked proteins (fibrinogen, fibronectin, hemoglobin and ferritin) were not cleaved (Fig. 2A,B; SI appendix, Fig. S5). PepOs from other bacteria may participate in the bacteria adhesion to and invasion of host cells through binding to human proteins such as plasminogen or fibronectin (Agarwal et al., 2013). However, PgPepO and TfPepO did not bind to any proteins they did not digest (SI appendix, Fig. S6). The hydrolysis sites of the cleaved substrates were identified using mass spectrometry and marked on the primary sequence of the substrates (Fig. 2C). We identified 6 and 9 unique hydrolysis sites for PgPepO and TfPepO, respectively, and seven sites shared by both enzymes. PgPepO cleaved the substrates near both terminuses of the substrates, a maximum of 8 amino acids away from it. On the contrary, TfPepO hydrolyzed α-casein deep inside the polypeptide chain – at least 50 aa from the terminuses of the protein, indicating endopeptidase character of this enzyme (Fig. 2C). To identify the motif recognized and hydrolyzed by PgPepO and TfPepO cleavage logos were generated (Fig. 2D). PgPepO and TfPepO were not characterized by restricted substrate specificity and only at the carbonyl side of the scissile peptide bond (P1’) large hydrophobic amino acids are preferred (Fig. 2D).

**Figure 2.**
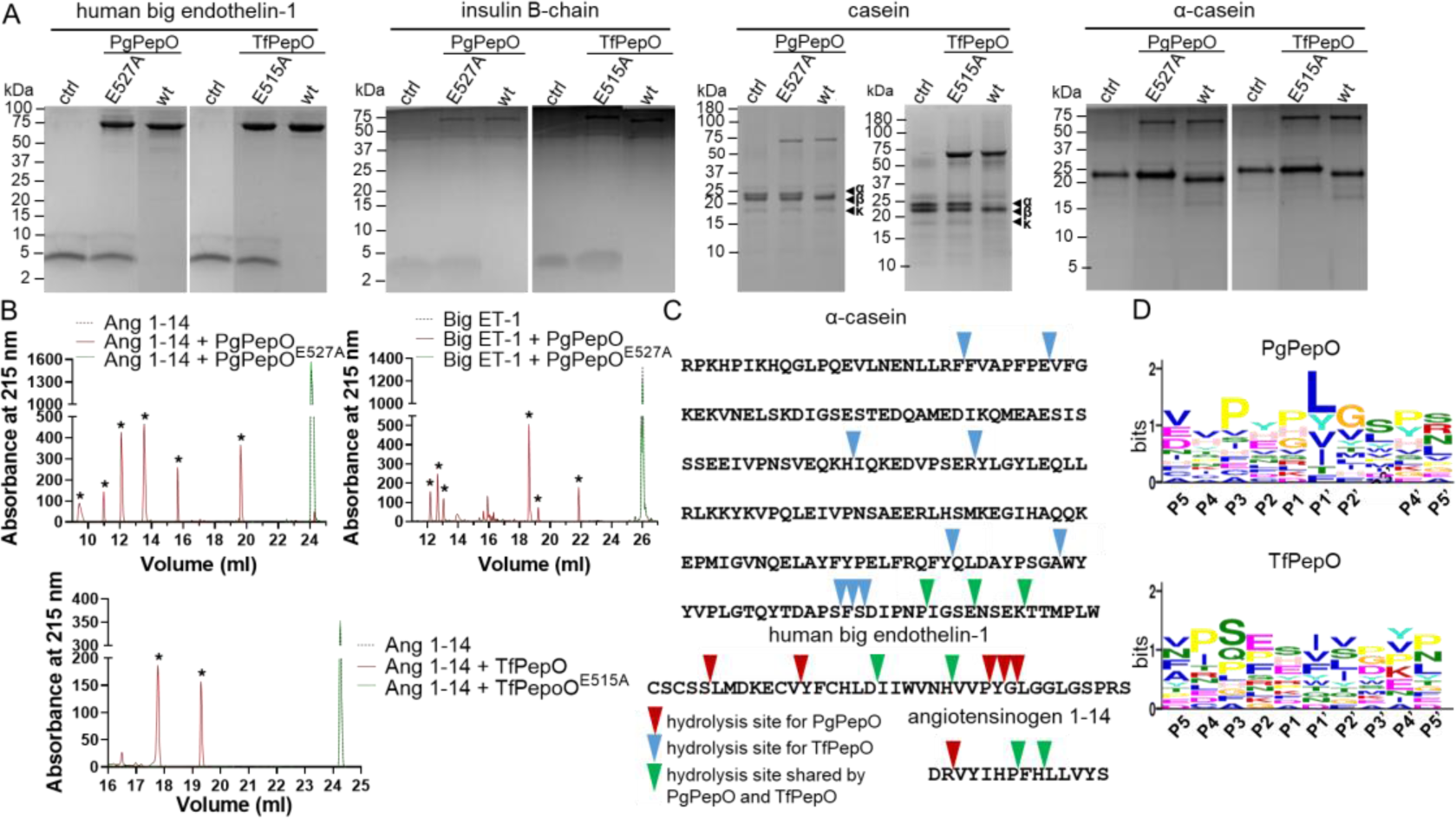
Determination of substrate specificity of PgPepO and TfPepO. (A) Different peptide and proteinaceous substrates were incubated alone (control, ctrl) and with PepOs: wild-type (wt) and catalytically inactive ExxxA mutants serving as negative controls, at substrate:enzyme weight ratios: 100:1 (big endothelin-1), 2000:1 (insulin B-chain), 80:1 (casein) and 200:1 (α-casein). Obtained samples were analysed by SDS-PAGE and mass spectrometry (MS). (B) Mixtures of the angiotensinogen 1-14 and human big endothelin-1 (Big ET-1) incubated at 800:1 substrate:enzyme weight ratios with PepOs: wt and E527A (PepO) or E515A (TfPepO) mutants were separated by High Performance Liquid Chromatography (HPLC). The content of peaks indicated by asterisks (*) were identified by MS (C, D) The identified by mass spectrometry cleavage sites for both PepOs were marked on the primary structures of the used substrates with differently colored triangles (C) and were used to create the cleavage logo employing Multiple Em for Motif Elicitation (MEME) (https://meme-suite.org) (D).

### Crystal structures of PgPepO and TfPepO

The solved crystal structures of PgPepO and TfPepO included variants missing short fragments, 14 (PgPepO) and 10 (TfPepO) residues, located at the N-terminus of the proteins. The secondary structure of PgPepO and TfPepO was mainly alpha helical in nature and could be further separated into two large lobe-like subdomains connected by intertwining polypeptide segments forming a smaller alpha helical linker region. Together with linker region both domains form a central, spherical cavity, which bears the active site of the proteases. The catalytic site of PepOs, located within the cavity surface of catalytic domain, was based around a zinc ion and conserved active site motif HEXXH (Fig. 3A,D). The catalytic zinc cation of PepOs was coordinated by the side chains of two histidines and glutamic acid (His526, His530, Glu575 for PgPepO; His514, His518, Glu575 for TfPepO) with the fourth coordination spot occupied by a water molecule. The proper position of histidines is ensured by interactions with two aspartic acids (Asp533 and Asp591 for PgPepO; Asp521, Asp579 for TfPepO). The zinc ion coordination sphere is completed by general base/acid glutamate (Glu527 and Glu515 in PgPepO and TfPepO, respectively), which polarizes the solvent and enhances its nucleophilicity (Fig. 3 B,E). Additionally, TfPepO contains second, structural zinc coordinated by Asp501, Asp 502, Lys602 and Arg610. Localization within the linker region suggested that TfPepO second, structural, zinc stabilize linker region, which was 25% longer than in PepO. The coordinated zinc ion and surrounding residues created a small binding pocket with subsites S1, S1′ and S2′ (Fig. 3C, F). Subsites S1 and S2’ are shallow and S1’ is deep, and its hydrophobic bottom is formed by the leucine side chain: 651 (PgPepO) and 639 (TfPepO). The shape and properties of S1’ explain the preference for hydrophobic amino acids at the P1’ position of both studied PepOs (Fig. 2D).

**Figure 3:**
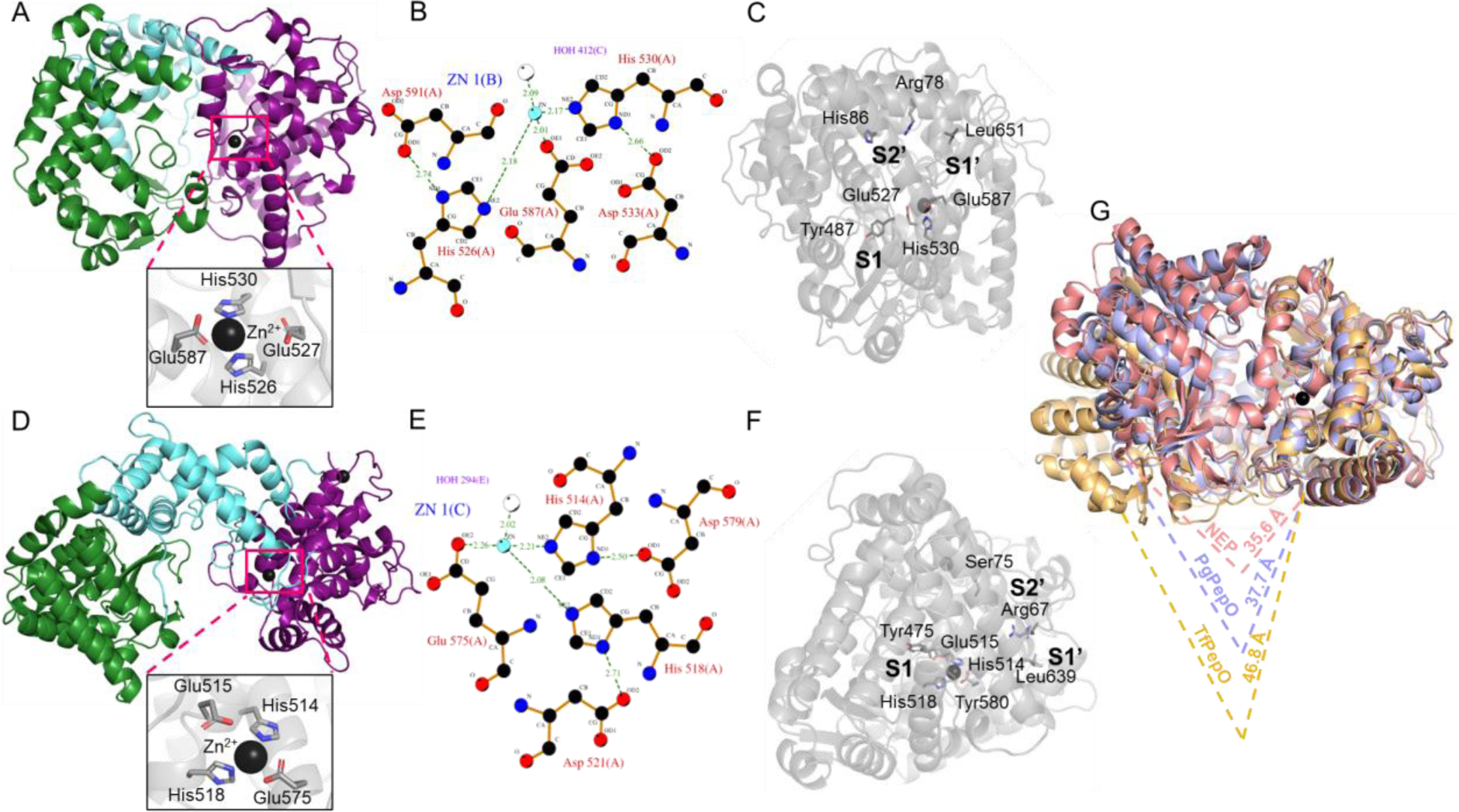
Crystal structures of PepOs. (A, D) General structure of PgPepO (A) and TfPepO (D) with a close-up of the zinc-binding site. (B, E) 2D diagram of interactions in the active site of PgPepO (B) and TfPepO (E) generated by PDBsum. C,F) PgPepO (C) and TfPepO (F) active site binding pocket labelled with subsites and residues. (E) Alignment of the structures of the PgPepO, TfPepO and neprilysin (PDB accession no: 6SH1), along with the width of the entrance to the catalytic site gap. All graphs and measurement were made using the PyMol program.

To better visualize the differences between PgPepO and TfPepO, a structural alignment with the structure of human neprilysin (PDB accession no.: 1DMT) was performed (Fig. 3G). The catalytic domains of all three M13 metalloproteases are characterized by low RMSD value (0.726-0.811) and higher differences can be observed in the linker regions and regulatory domains as well as the arrangement of the regulatory domains relative to the active sites. The regulatory domain of TfPePO is further away from the catalytic domain, and thus the space between both domains is larger than in the case of PgPePO and neprilysin. Described observations were confirmed by measuring the width of the entrance to the catalytic site cleft. For PgPepO and neprilysin it was very similar and equal to 37.7 and 35.6 Å, respectively, and for TfPepO it was over 9 Å longer - 46.8 Å (Fig. 3G).

### PepOs are attached to the cell surface

At the N-terminus both PgPepO and TfPepO contain a putative signal peptide characteristic for secretory lipoproteins (SI appendix, Fig. S1). Upon its removal during translocation across the inner membrane, the conserved cysteine becomes the novel N-terminus of the protein. Then, this N-terminal Cys is modified by attaching a lipid providing the membrane anchorage. It is however, impossible to predict whether the lipoprotein will remain in the inner membrane or be transported to the outer membrane, based on the protein sequence alone. In the latter case, the lipoprotein may be located in the outer membrane inner leaflet or the outer membrane outer leaflet. We used anti-PepO antibodies to detect the proteins of interest in the fractions obtained from *P. gingivalis* and *T. forsythia* cultures. PgPepO and TfPepO were located in the cell envelope fraction containing both cell membranes and peptidoglycan, as well as in the outer membrane vesicles (OMV) (Fig. 4A). This observation indicates that PepOs are attached to the cell surface. To confirm it, we performed dot-blot analysis, in which inner membrane biotinylated proteins (control of cell integrity) and PepO deficient mutant strains (ΔPepO) were used as a control. The identical signal in both whole cells and lysed cells confirmed that the PepOs are attached to the *P. gingivalis* and *T. forsythia* cell surface (Fig. 4B). This conclusion was clearly confirmed by flow cytometry analysis (Fig. 4C, SI appendix, Fig. S7) and, by confocal microscopy (only for TfPepO; SI appendix, Fig. S8). The findings are further corroborated by mass spectrometry analysis showing the presence of PgPepO (Veith et al., 2014) and TfPepO (Friedrich et al., 2015) in OMV and, only for *T. forsythia*, outer membrane (Veith et al., 2009). Next, we checked whether whole cultures of *P. gingivalis* and *T. forsythia* or their purified proteinases were active against the substrate Mca-RPPGFSAFK(Dnp)-OH and our results indicate that whole cultures were highly efficient in the substrate processing, while previously described proteases of both bacteria were not responsible for this activity (SI appendix, Fig. S9). To deepen it, we measured the activity towards the substrate in fractions derived from the culture of both bacteria: wild-type and PepO deletion variants. The proteolytic activity of whole cultures was split between cells and OMVs. The use of deletion mutants showed that PepO was responsible for all measured activity in *T. forsythia*. However, in *P. gingivalis*, PepO deletion only statistically significantly reduced the activity by 42% (cells, C) and 29% (OMV) (Fig. 4D). This suggests that, unlike *T. forsythia*, *P. gingivalis* besides PepO secretes at least one additional protease capable of hydrolyzing the substrate.

**Figure 4.**
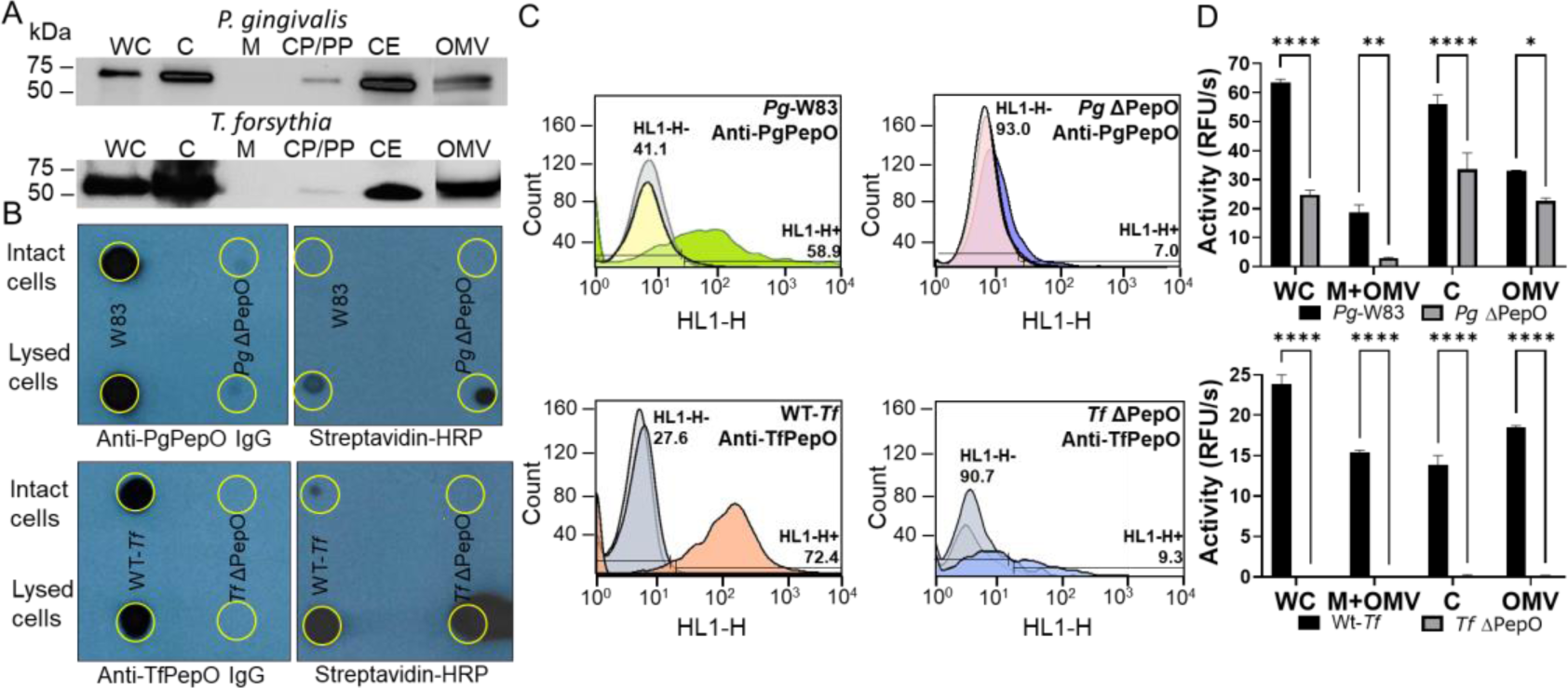
PgPepO and TfPepO are attached to the cell surface. (A) Western blot of culture fractions: whole culture (WC), cells (C), particle-free culture medium (M), soluble proteins derived from cytoplasm and periplasm (CP/PP), cell envelope (CE) and outer-membrane vesicles (OMV) isolated from *P. gingivalis* and *T. forsythia* cultures. Blots were probed with rabbit polyclonal antibodies anti-PepO. (B) Dot-blot analysis of *P. gingivalis*: W83 (wild-type, wt) and *Pg* ΔPepO (upper panels) and *T. forsythia*: wt (WT-*Tf*) and *Tf* ΔPepO (lower panels) intact (upper row) and lysed by sonication (lower row) cells to detect PepOs (left panels) and biotinylated inner membrane (IM) protein (right panels). Dots placed on the PVDF membrane (dotted circle) were probed with rabbit polyclonal antibodies anti-PepOs and streptavidin conjugated with HRP (streptavidin-HRP). (C) Flow cytometry analysis showing the surface exposure of PepOs in *P. gingivalis* (upper histograms): W83 (wt) and ΔPepO, and *T. forsythia (*lower histograms): wt (WT-*Tf*) and ΔPepO using appropriate anti-PepOs antibodies. Brighter colors on histograms represent negative controls: cells alone or stained with secondary antibodies. (D) Comparison of activity against Mca-RPPGFSAFK(Dnp)-OH in obtained culture fractions: WC, C, cell-free culture medium (M+OMV) and outer-membrane vesicles (OMV) derived from *P. gingivalis*: W83 and ΔPepO (upper graph) and *T. forsythia*: wt (WT-*Tf*) and ΔPepO (lower graph). Statistical differences were determined by one-way ANOVA (*P<0.05; **P<0.01; ****P<0.0001).

### PepOs are proteases responsible for Ang I cleavage

*P. gingivalis* and *T. forsythia* secrete numerous proteases. To exclude the possibility that proteases other than PepO hydrolyze Ang I, this peptide was incubated with a cell-free medium from the culture of both periodontopathogens, which contains both proteases secreted directly into the medium (e.g. *T. forsythia* KLIKK proteases) and those bound to the cell surface and released in the form of OMV (e.g. *P. gingivalis* gingipains). In both cases, Ang I was hydrolyzed to Ang 1-9. Ang 1-7 could not be clearly identified due to the separation at the same elution time peptides probably derived from components of the medium used. The lack of Ang I degradation by PepO-deficient mutants indicates that PepOs were the only proteases responsible for Ang I hydrolysis by *P. gingivalis* and *T. forsythia* (Fig. 5A).

**Figure 5.**
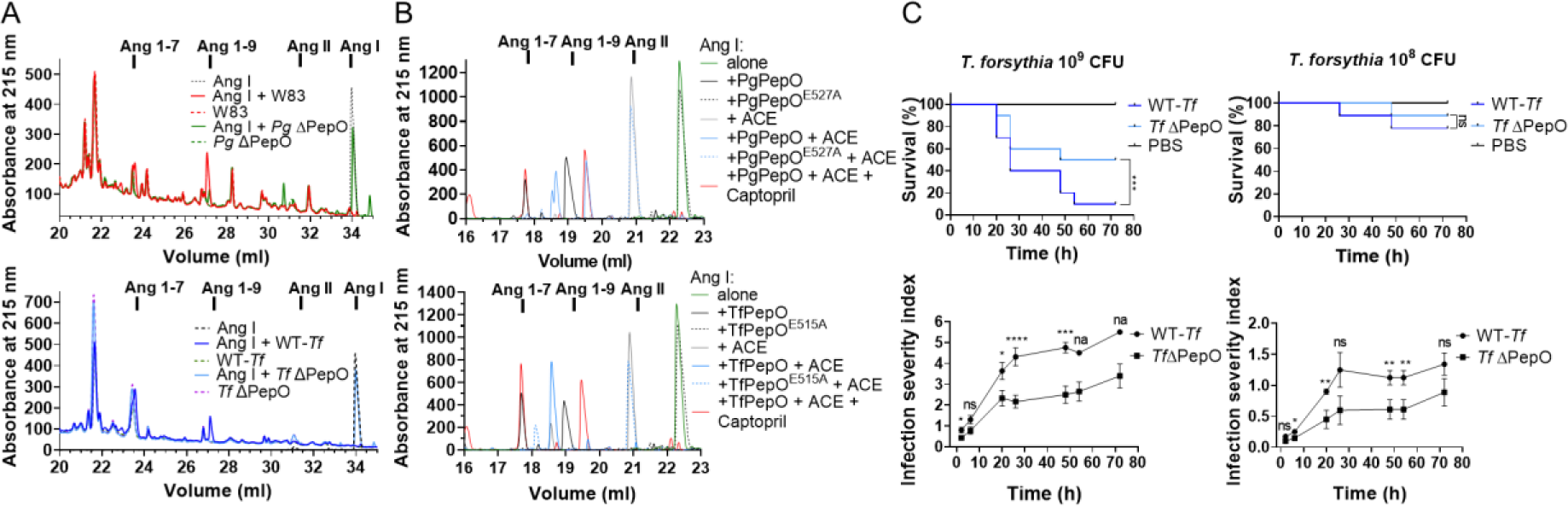
PepOs are responsible for the cleavage of Ang I by *T. forsythia* and *P. gingivalis* and contribute to *T. forsythia* virulence *in vivo*. (A) Ang I was incubated with cell-free medium (M+OMV) derived from *P. gingivalis*: W83 and ΔPepO (upper row) or *T. forsythia*: wild-type (WT-Tf) and ΔPepO (lower row) and then separated on HPLC column. Products of Ang I proteolysis were identified based on pure angiotensins separated under the same conditions. (B) Ang I was incubated with different enzymes, PgPepO, TfPepO, and their inactive mutants, as well as ACE in various configurations, and then separated on the HPLC column. Products of Ang I proteolysis were identified based on pure angiotensins separated under the same conditions. (C) *Galleria mellonella* larvae (N=10 for each group) were infected with two different doses of *T. forsythia*: WT-Tf and *Tf*ΔPepO. Dead and live larvae were counted at indicated time points, and the infection severity index was calculated based on the assessment of mobility and melanisation of larvae. The data are mean±SD and are representative of three biological replicates (C). Statistical differences were determined by one-way ANOVA or Kaplan-Meier (survival curves) test (*P < 0.05; **P < 0.01; ***P < 0.001; ****P < 0.0001; na - not available - single larvae alive).

ACE is the primary enzyme that directs RAS to form Ang II in people. On the contrary, the PgPepO and TfPepO were responsible for forming Ang 1-9 and Ang 1-7. Therefore, we checked what product from Ang I is formed in the presence of an equimolar mixture of ACE and PepOs (Fig. 5B). As expected, ACE alone generated Ang II. However, the addition of PgPepO or TfPepO resulted in the formation of no active peptide: neither Ang II nor Ang 1-7. These results may be explained by the ability of ACE to hydrolyze Ang 1-7 (Deddish et al., 1998). However, adding captopril, an ACE inhibitor used to treat hypertension, to the mixture resulted in the formation of only Ang 1-7, with a complete lack of Ang II (Fig. 5B).

### PepO contributes to *T. forsythia* virulence in the *Galleria mellonella* model

*P. gingivalis* and *T. forsythia* are asaccharolytic bacteria that use peptides, not sugars, as an energy source. Therefore, it can be assumed that PepO, together with other proteases, are involved in the degradation of proteins into short peptides, which can then be used in the metabolism of both bacteria. To verify it, we checked the effect of PepO deletion on the growth of *P. gingivalis* and *T. forsythia* in a minimal medium consisting of DMEM with 2% BSA. The lack of PepO practically did not affect the growth of both bacteria, indicating that PepOs do not participate in peptide acquisition by both pathogens (SI appendix, Figure S10).

The immune system of *G. mellonella* includes complement-like proteins, opsonins, phagocytic cells (hemocytes), and antimicrobial peptides, thus showing significant similarities to the human innate immune system (Ménard et al., 2021). This model was used to show that *Streptococcus mutans* PepO contributes to the virulence of this bacterium (Alves et al., 2023). Thus, *G. mellonella* larvae were infected with *T. forsythia* and *P. gingivalis*: wild-type strain (WT or W83) and a mutant lacking PepO (ΔPepO) in two doses: 10^8^ and 10^9^ CFU. Then, the survivability and the severity of infection, assessed based on melanization and larval activity, were monitored. In the case of *P. gingivalis*, no differences were observed between the wild-type strain and the deletion mutant (SI appendix, Figure S11). On the contrary, the lack of PepO in *T. forsythia* resulted in higher larvae survival (only for 10^9^ dose) and weaker infection symptoms (Fig. 5C).

## Discussion

The renin-angiotensin system, mainly responsible for regulating blood pressure, is also active in different tissues. Moreover, the correlation between high blood pressure and periodontitis is described and taking ACE inhibitors for hypertension worsens periodontal health. In this study, we show that homologous surface attached proteases of two periodontopathogens, *P. gingivalis* (PgPepO) and *T. forsythia* (TfPepO), have an oppositeactivity to ACE because they process Ang I not into Ang II, but Ang 1-7.

TfPepO and PgPepO share substrate specificity, i.e., a preference for large hydrophobic amino acids at the carbonyl side of the scissile peptide bond, with other human and bacterial members of the M13 family: neprilysin, ECE1, ECE2, *Mycobacterium tuberculosis* Zmp-1 and *S. pneumoniae* PepO, and ACE2 (M2 family) (Rawlings et al., 2018). Despite it and the high homology within the catalytic domain, the TfPepO and PgPepO are not inhibited by any tested inhibitor (Fig. 1F). The answer to this is provided by the analysis of the crystal structure of neprilysin in complex with the inhibitor LBQ657 (PDB accession no: 5JMY). (Schiering et al., 2016). This compound occupies the S1, S1’ and S2’ subsites within the catalytic cleft. The methyl group of LBQ657 forms a hydrophobic bond with Phe544 forming the shallow S1 pocket, and P1’ biphenyl binds to the deep S1’ pocket. Additionally, P2’ succinic acid interacts with Arg102 and Arg110. In the case of TfPepO and PgPepO, interaction only with the S1’ pocket is possible because there is Tyr instead of Phe in S1, and only one of the two Arg positioning the carboxylic acid in the S2’ subsite is preserved in PgPepO and TfPepO structures (Fig. 3C,F). These differences probably explain the lack of inhibition of the tested PepOs by LBQ657.

TfPepO, unlike PgPepO and other M13 metalloproteases, hydrolyzed peptide bonds at a much greater distance from the terminuses of the substrates (Fig. 2C). The explanation for this distinctive feature of TfPepO is the much wider entrance to the catalytic center, which allows bigger substrates to access it. It is probably possible due to the more extended linker region in TfPepO (25% and 35% longer than in PgPepO and neprilysin, respectively), stabilized by additional structural zinc (absent in other members of the M13 family of proteases), which provides greater separation between the catalytic and regulatory domain. Therefore, the width of the entrance to the catalytic cleft was also measured in other members of the M13 family of metalloproteases (SI appendix, Fig. S12). In all cases, the width is similar to PgPepO and neprilysin and is 30.7-37.7 Å, and is significantly shorter than in TfPepO (46.8 Å). The wider entrance to the catalytic cleft is probably a unique feature of TfPepO that distinguishes it from other members of the M13 family.

After removal of the signal peptide during translocation through the inner membrane, PgPepO and TfPepO still have 19 and 17 amino acid residues long, respectively, unstructured fragments preceding the molecule core (Fig. 3A). This fragment probably ensures appropriate separation from the membrane, thus enabling unrestricted activity of PepO on the bacterial surface. The *T. forsythia* cell is additionally surrounded by a proteinaceous crystalline S-layer, which can limit access of substrates to TfPepO by physically blocking the access of substrates to the enzyme. However, the detection of TfPepO using IgG antibodies in flow cytometry indicates that the S-layer does not limit the activity of TfPepO in any way. This conclusion is additionally confirmed by the fact that another *T. forsythia* surface lipoprotein, serpin miropin, despite the presence of the S-layer, inhibits even human plasmin (∼80 kDa) through the formation of a covalent complex (Sochaj-Gregorczyk et al., 2020).

The biological function of PepO has been described in other bacteria. PepO protects *Streptococcus sanguinis* and *Streptococcus pyogenes* from bactericidal activity of the complement system (Honda-Ogawa et al., 2017) and, in the former case, also increases survival in the *Galleria mellonella* model (Alves et al., 2023). *Streptococcus pneumoniae* PepO binds to plasminogen and fibronectin and participates in immune evasion and host cell invasion (Agarwal et al., 2013; Agarwal et al., 2014).

*M. tuberculosis* PepO, Zmp-1 increases the pathogen’s survival inside macrophages by blocking the inflammasome activation (Master et al., 2008; Paolino et al., 2018). The results of *G. mellonella* model showed that TfPepO is probably involved in the evasion of host defence mechanisms (Fig. 5), and PgPepO is only responsible for host cell invasion (Ansai et al., 2002; Miller et al., 2017). The different functions of the studied PepOs can be explained by the fact that in *P. gingivalis* only gingipains, ∼85% of overall proteolytic activity, are responsible for avoiding the host’s response (Imamura 2003; Potempa & Pike, 2005; Guo et al., 2010; Bengtsson et al., 2015), and in *T. forsythia* these roles are distributed among more proteases, e.g. KLIKK proteases and TfPepO (Karim et al., 2010; Koziel et al., 2010; Koneru et al., 2017).

The RAS system plays an important role not only in the regulation of the circulatory system and kidney function but is also active in periodontal tissues (Saravi et al., 2020), where the expression or presence of all critical elements of this system, i.e. angiotensinogen, renin, ACE, ACE-2, Ang I, Ang 1-9, Ang II, Ang 1-7, AT1 and AT2 receptors (Nakamura et al., 2011), as well as the Mas receptor (SI appendix, Fig. S13). The formation of Ang II, in addition to increasing blood pressure, increases inflammation and oxidative stress (Weber & Brilla, 1991; Welch 2008). Moreover, reduced Ang II signaling result in bone loss in mice (Lima et al., 2021). PgPepO and TfPepO lead to the formation of Ang 1-7 from Ang I, and in the presence of ACE, they prevent the formation of Ang II, so they probably reduce the level of reactive oxygen species at the site of infection. It may be crucial for *P. gingivalis* and *T. forsythia* obligate anaerobes and protection of their virulence factors against oxidation (Fitzsimonds et al., 2021). The proposed anti-inflammatory role of PgPepO and TfPepO is partially supported by the observation that the deletion of neprilysin, responsible for the metabolism of proinflammatory peptides, increases the sensitivity of mice to endotoxic shock (Lu et al., 1995).

Many clinical studies describe a greater risk of hypertension in people suffering from periodontitis. Curing periodontitis does not significantly decrease blood pressure (Muñoz Aguilera et al., 2020). However, people taking ACE inhibitors to treat hypertension have worse periodontal health (Rodrigues et al., 2016; Wang et al., 2020; Chatzopoulos et al., 2023; Frankenhaeuser et al., 2023). Given that in the presence of an ACE and its inhibitor, captopril, PgPepO and TfPepO not only degrade ACE-generated Ang II but produce Ang 1-7 from Ang I (Fig. 5B), it is tempting to postulate that the activity of bacterial PepO contributes to the observed deterioration of periodontal health in people taking ACE inhibitors. While it is premature to state that PgPepO and TfPepO activity contributes to periodontitis progression in such patients, our data demonstrate that among numerous proteases secreted by *P. gingivali*s and *T. forsythia,* PgPepO and TfPepO, respectively, can interfere with RAS system through Ang I cleavage.

## Materials and methods

### Reagents

Fast Digest BamHI and EcoRI, dNTP, GeneJET™ Gel Extraction Kit, GeneJET™ PCR Purification Kit, GeneJET™ Plasmid Miniprep Kit, GeneRuler 1 kb DNA Ladder, GeneRuler 50 bp DNA Ladder, Pierce™ ECL Western Blotting Substrate, Vials with glass insert, Micro+™, Chromacol™, Goat anti-Rabbit IgG Alexa Fluor™ 488, Hoechst 33342 were acquired from Thermo Scientific (Waltham, MA, USA). The QuikChange Lightning Site-Directed Mutagenesis Kit was purchased from Agilent Technologies (Santa Clara, CA, USA). All primers used in the study were synthesized by Genomed (Warsaw, Poland). T4 DNA ligase, Platinum SuperFi DNA Polymerase, Streptavidin-HRP, Streptavidin-FITC, were purchased from Invitrogen (Waltham, Massachusetts, USA). Genomic Mini System was from A&A Biotechnology (Gdańsk, Poland). The expression vector pGEX-6P-1, glutathione-Sepharose 4 Fast Flow, HiLoad 16/600 Superdex 200 pg and Superdex 200 increase 10/300 GL columns, LMW and HMW protein kit were obtained from GE Healthcare Life Sciences (Little Chalfont, UK). Aeris C18: analytical column, Aeris 3.6 μm PEPTIDE XB-C18 100 Å, LC Column 150 × 4,6 mm was from Phenomenex (Torrance, CA, USA). Tryptic soy broth (TSB), hemin, N-acetylmuramic acid (NAM), Menadione, E-64, Leupeptin, Captopril, Lisinopril, Orthophenantroline, 1,10-Phenantroline (OF), Pefabloc, Phosphoramidon, Sacubitrilat (LBQ657), Tiorphan, α-casein, angiotensin I, angiotensin II, angiotensin 1-7, angiotensinogen 1-14, human Big endothelin-1, endothelin 1, fibrinogen, fibronectin, hemoglobin, casein, LL-37, insulin β-chain, Mca-(Ala7,Lys(Dnp)9)-bradykinin (Mca-RPPGFSAFK(Dnp), Peroxidase-conjugated Anti-Rabbit IgG (whole molecule) and concentrators Amicon Ultra-15 Centrifugal Filters, 10k Da MWCO, 15 mL were purchased from Merck (Darmstadt, Germany). Fetal bovine serum (FBS) was from Gibco (Dublin, Ireland) and defibrinated sheep blood was from Graso Biotech (Starogard Gdański, Poland). Daglutril (SLV-306) was purchases from Axon Medchem (Groningen, Holland), FP-biotin was from Santa Cruz Biotechnology, Inc. (Dallas, TX, USA), Human [D-Val22]-Big Endothelin-1 (16-38) was acquired from Biomatik (Kitchener, Ontario, Canada), Iodoacetamid (IAA) from VWR, (Radnor, PA, USA) and angiotensin 1-7 was from APExBIO (Houston, TX, USA). Polyclonal rabbit primary antibodies against PgPepO and TfPepO were obtained from Proteogenix (Schiltigheim, France). The protein molecular weight standards Precision Plus Protein™ Dual Xtra Prestained Protein Standards and Precision Plus Protein™ Kaleidoscope™ Prestained Protein Standards, Polyvinylidene difluoride (PVDF) and nitrocellulose membranes were purchased from Bio-Rad (Hercules, CA, USA). Complete EDTA-free protease inhibitor cocktail was from Hoffmann-La Roche (Basel, Switzerland). Syringe filters (0.22 and 0.45 μm) and 96-well transparent plates were purchased from Sarstedt (Blizne Łaszczyńskiego, Poland), Medical X-Ray-Film Blue (X-ray films) from AGFA (Mortsel, Belgium) and 96-well black plates were from Nunc (Roskilde, Denmark). *Galleria mellonella* larvae were acquired from Egzotic Room (Plewiska, Poland). All other chemical reagents were obtained from BioShop Canada (Burlington, ON, Canada).

### Bacterial growth and culture fractionation

*T. forsythia* ATCC 43037: wild-type (WT) and *Tf* ΔPepO, and *P. gingivalis*: W83 (WT) and *Pg* ΔPepO were grown in an anaerobic chamber (Whitley A85: Don Whitley Scientific, Bingley, Great Britain) in an atmosphere of 90% nitrogen, 5% carbon dioxide and 5% hydrogen at 37°C. Liquid cultures of both bacteria were grown in enriched TSB (eTSB) broth containing 30 g/l TSB broth and 5 g/l yeast extract, supplemented with 5 mg/l hemin, 0.5 g/l L-cysteine and 2 mg/l menadione for *P. gingivalis* (PgeTSB), 5% fetal bovine serum and 10 mg/l N-acetylmuramic acid for *T. forsythia* (TfeTSB). The solid medium for *P. gingivalis* and *T. forsythia* contained the appropriate eTSB medium with 1.5% agar and 5% defibrinated sheep blood (Graso Biotech, Starogard Gdański, Poland). *Tf* ΔPepO and *Pg* ΔPepO were additionally grown in the presence of 5 μg/ml erythromycin. Bacterial cultures of *T. forsythia* (WT and *Tf* ΔPepO) with an optical density (OD_600 nm_) = 0.4 - 0.6 and *P. gingivalis* (W83 and *Pg* ΔPepO) with OD_600 nm_ = 1.2 - 1.5 were adjusted to OD_600 nm_ = 1.0 with PBS. The adjusted cultures (whole culture fraction, WC) were centrifuged (10 min, 4 500 × g, 4°C), the supernatant (cell-free medium) was collected, and the cells (cells fraction, C) were suspended in PBS alone or supplemented, only for samples for immunoassays, with complete EDTA-free protease inhibitor cocktail (Hoffmann-La Roche, Basel, Switzerland) and 200 µM TLCK (only for *P. gingivalis*). The supernatant was filtered (0.45 μm) and ultracentrifuged (1 h, 150 000 × g, 4°C). The particles-free supernatant (medium fraction, M) was collected, and the pellet was washed with 25 ml of fractionation buffer (FB): 20 mM Tris, 150 mM NaCl, 5 mM MgCl_2_, 0,02% (w/v) NaN_3_, (pH 7.5). Ultracentrifugation was repeated, and the obtained pellet (outer membrane vesicles fraction, OMV) was suspended in 250 μl of PBS with, only for samples for immunoassays, the addition of appropriate inhibitors. The cells were disrupted by ultrasound employing UP50H ultrasonic homogenizer (Hielscher Ultrasonics GmbH, Teltow, Germany): 10 cycles of 10 pulses of 0.5 sec each, pulse amplitude 60%, then centrifuged (20 min, 10 000 × g, 4°C) and the obtained supernatant was ultracentrifuged (1 h, 150 000 × g, 4°C). The supernatant (cytoplasm/periplasm fraction, CP/PP) was collected, and the pellet was washed with 25 ml of FB. Ultracentrifugation was repeated, and the pellet was suspended in PBS alone or only for samples for immunoassays, with the addition of appropriate inhibitors (cell envelope fraction, CE).

### Cloning and mutant construction

The coding sequence of the full-length PgPepO gene (KEGG accession number: K07386) without the predicted signal peptide (SP) (^23^C-^689^W) was amplified by PCR directly from genomic DNA isolated from *P. gingivalis* W83 using the primers listed in Table S1. Next, the pGEX-6P-1 vector and the PCR product were digested with BamHI/EcoRI and ligated. The sequence encoding TfPepO without SP, (^17^C-^677^W), (GenBank accession number: KKY61269.1) was synthesized and cloned into pGEX-6P-1 using BamHI/XhoI by Genscript (Nanjing, China). The wild-type (WT) constructs (pGEX-6P-1-PgPepO, pGEX-6P-1-TfPepO) were used to obtain catalytically inactive mutants (PgPepO^E527A^ and TfPepO^E515A^) in which the glutamic acid in the active site was substituted by alanine employing QuikChange Lightning Site-Directed Mutagenesis Kit (Agilent Technologies) and appropriate primers listed in SI appendix, Table S1. The correctness of genetic constructs was confirmed by DNA sequencing (Genomed, Warsaw, Poland).

### Enzymatic activity assays

PepOs activity, unless otherwise mentioned, was determined in 100 μl in reaction buffer, 100 mM Tris, 150 mM NaCl, 2.5 mM CaCl_2_, 0.02% NaN_3_, 0.05% Pluronic F-127 (pH 7.5), using Mca-RPPGFSAFK(Dnp)-OH as a substrate. The enzymatic reaction was monitored in the wells of a 96-well black plate at 37°C as an increase in fluorescence (excitation/emission = 320/405 nm) over time using a SpectraMax Gemini EM reader (Molecular Devices, San Jose, CA, USA). The Michaelis-Menten constant (K_m_) was calculated using the GraphPad Prism v9 based on reaction velocity (V) measured at different concentrations of substrate (1-50 μM). For the pH optimum, PepOs activity was determined using the buffer: 100 mM Tris, 50 mM MES, 50 mM acetic acid, 0.02% NaN_3_ at different pH ranging from 5.0 to 11.0. For the temperature optimum, activity was measured at different temperatures ranging from 4 to 99°C for 30 min and enzymatic reaction was stopped by addition of 20 mM OF. To test the effect of temperature, inhibitors, reducing agents, and divalent metal cations on PepOs activity, the proteases were incubated in temperatures (4-99°C) in reaction buffer for 30 min or with each additive in 100 mM Tris, 0.02% NaN_3_ (pH 7.5) for 15 min at 37°C and the enzymatic activity was determined as described above.

### Determination of PepOs cleavage specificity

Proteins: human albumin, ferritin, fibrinogen, fibronectin, haemoglobin, bovine casein, α-casein, and the peptides: human big endothelin-1, human cathelicidin LL-37, bovine insulin B-chain (4 μg) were incubated with PgPepO and TfPepO at the enzyme/substrate weight ratios 1:100 (big endothelin-1), 1:200 (α-casein, LL-37), 1:2000 (insulin B-chain), and 1:1000 (LL-37) and 1:80 (other proteins) for 24 h (α-casein, big endothelin-1, insulin β-chain) or seven days (other substrates) at 37°C in 100 mM Tris, 150 mM NaCl, 2.5 mM CaCl_2_, 0.02% NaN_3_, 0.05% Pluronic F-127 (pH 7.5). Obtained samples were analyzed by SDS-PAGE (Schägger & Jagow, 1987): 4/12% and 4/18% (T:C, 27.5:1) gels for proteinaceous and peptide substrates, respectively, and by mass spectrometry. Substrates incubated with PgPepO^E527A^ and TfPepO^E515A^ were used as a negative control.

Angiotensinogen 1-14 and human big endothelin-1 (30 µg) were mixed with PepOs (final concentration: 10 nM) and incubated 2 h at 37°C. Then, trichloroacetic acid (TFA) was added (final concentration 0.1%). Obtained samples were centrifuged (10 min, 16,900 × g, 4°C), and the supernatant was separated on an Aeris 3.6µm Peptide XB-C18 column (150 × 4.6 mm) in an acetonitrile gradient (2-60%). Samples containing substrate: alone and with an inactive PepO mutant were used as controls. The Mass Spectrometry Laboratory (Institute of Biochemistry and Biophysics, Polish Academy of Sciences, Warsaw, Poland) identified proteolysis products.

### Deletion of PepO proteases in *P. gingivalis* and *T. forsythia* bacteria

DNA fragments containing erythromycin resistance cassette (GenBank accession number: AF219231.1) (Fletcher *et al*. 1995) flanked by upstream and downstream fragments surrounding PepOs gene in genomes were synthesized and cloned into pUC19 plasmid by Genscript (Nanjing, China). Deletion mutants were obtained by natural homogenous recombination. 1 µg of suicide vector and 100 µl of electrocompetent cells were transferred in a sterile, ice-cooled electroporation cuvette (Bio-Rad), which was placed in the MicroPulser^TM^ electroporator (Bio-Rad) and subjected to an electric pulse: 2.5 kV, 3 msec. Immediately after electroporation, 1 ml of the appropriate PgeTSB or TfeTSB medium preheated at 37°C was added and obtained mixtures were kept overnight (∼16-24 h) at 37°C under anaerobic conditions. Then, the bacterial mixture was spread on selective plates with the addition of erythromycin (5 µg/ml). Genomic integration was confirmed by amplifying genomic fragments containing introduced mutations and sequencing them (Table S1).

### Western blot, dot blot and flow cytometry analysis

For Western blot, resolved proteins by SDS-PAGE were electrotransferred (1 h, 90 V, 10 mM CAPS, 10% (v/v) methanol (pH 11.0)) onto PVDF membranes. Membranes were blocked in EveryBlot Blocking Buffer (BioR) for 30 min, then probed with the appropriate polyclonal rabbit antibody against PepOs at concentration 1μg/ml in EveryBlot Blocking Buffer (Bio-Rad) followed by goat anti-rabbit IgG horseradish peroxidase-conjugated antibodies (1:20000) (Merck), and in between membranes were washed in TTBS, 20 mM Tris, 150 mM NaCl, 0.02% (w/v) NaN_3_, 0.1% Tween-20 (pH 7.5). The blots were developed using the Pierce™ ECL Western Blotting Substrate (Thermo Scientific) substrate kit and X-ray (Agfa, Mortsel, Belgium) films or ChemiDoc™ Touch Imaging System (Bio-rad). For dot-blot analysis, isolated cells were divided into two parts. One part was supplemented with SDS (a final concentration of 0.05%) and disrupted by sonication (10 cycles, 10 pulses of 0.5 sec. each, pulse amplitude 60%) with a UP50H device (Hielscher Ultrasonics). 2.5 μl of intact and sonicated cells were spotted onto PVDF membranes and analyzed as described above. For flow cytometry, cells’ suspension from cultures adjusted to OD_600 nm_ = 1.0 was obtained similarly, with one exception: PBS-BSA buffer (PBS with the addition of 1.5% (w/v) BSA and complete EDTA-free protease inhibitor cocktail) was used. 100 μl of cell suspension was transferred to a 96-well conical plate, collected by centrifugation (1000 × g, 5 min, 4°C). The pellets were probed for 30 min on ice with 100 μl of 30 μg/ml polyclonal rabbit anti-PepO antibody in PBS-BSA, followed by incubation with 100 μl of goat anti-rabbit antibody conjugated with Alexa Fluor 488 at 1:200 dilution or probed only with Streptavidin−FITC at 1:200 dilution. After 30 min of incubation on ice, cells were washed three times with PBS-BSA and analyzed using a Guava® easyCyte™ flow cytometer (Luminex Corporation, Austin, TX, USA) and InCyte™ Software (Merck). Cells with no antibodies or only the secondary antibodies were used as controls.

### Analysis of Ang I hydrolysis by media after cultivation of *P. gingivalis* and *T. forsythia*

Bacterial cultures were centrifuged (10 min, 4500 × g, 4°C), and supernatant (cell-free medium, M+OMV fraction) was collected. Reaction mixtures (final volume: 50 µl) containing 30 µg of the following substrates: angiotensin I (Ang I) and an amount of supernatant with activity against Mca-RPPGFSAFK(Dnp)-OH equal to 10 nM of recombinant PepO. Samples were incubated for 72 h at 37°C, then trichloroacetic acid (TCA) was added to the final concentration of 3.5%, and samples were kept for 15 min on ice. TFA (final concentration: 0.1%) was added to the mixtures, and samples were centrifuged (10 min, 16 900 × g, 4° C). The supernatant was separated on an Aeris 3.6um Peptide XB-C18 column (150 x 4.6 mm) in an acetonitrile gradient (2-80%). As controls, samples containing only the substrate, substrate and supernatant from PepO deletion mutants, and appropriate eTSB medium.

### Virulence of P. gingivalis and T. forsythia in the Galleria mellonella model

Only healthy larvae (Egzotic Room, Plewiska, Poland) without signs of melanization were used in the experiments. Groups of 10 larvae were injected into the last pair of legs using a Hamilton syringe with 10 μl of a bacterial suspension containing 10^8^ or 10^9^ CFU of *P. gingivalis* (W83 and *Pg* ΔPepO) and *T. forsythia* (WT and *Tf* ΔPepO) cells. 10 µl of PBS was used as a control. Larvae were kept at 37°C in 9 cm Petri dishes without food. Their condition regarding physical activity and melanization was monitored for up to 72 h after infection and subjected to scoring. The experiment was performed in three biological replicates.

### Crystallization of protein

Purified proteins in 20 mM Tris, 50 mM NaCl, pH 8.0 were concentrated to 10 mg/ml. Screening for crystallization conditions was performed using commercially available buffer sets in a sitting-drop vapor diffusion setup by mixing 0.2 μl of protein complex solution and 0.2 μl of buffer solution. Crystals of PgPepO were obtained at room temperature from solutions containing Tris pH=7.5, 0.2 M magnesium chloride, 40% (v/v) EDO, 15% (w/v) PEG 8000 (pH=6.5), while crystals of TfPepO were obtained at room temperature from solutions containing 0.3 M sodium fluoride, 0.3 M sodium bromide, 0.3 M sodium iodide, 0.1M buffer system containing imidazole / MES mix, 40% (v/v) ethylene glycol, 20% (w/v) PEG 8000, pH 6.5.

### Structure Determination and Refinement

Crystals were flash-cooled in liquid nitrogen. The crystallographic experiments were performed on the 14.1 beamline at the HZB Bessy, Berlin, Germany (PgPepO) and the 11.2C beamline at the Elettra Synchrotron, Trieste, Italy (TfPepO).. The data were indexed, integrated and scaled using XDS (Kabsch 2010) and subsequently merged using Aimless (Evans & Murshudov, 2010). PgPepO structure was solved using AutoSol and anomalous data collected at zinc wavelength and resulting initial model was used for molecular replacement. TfPepO structure was solved by molecular replacement using MORDA pipeline and 3DWB PDB entry as a search model (Vagin & Lebedev, 2015; Schulz et al., 2009). Models was automatically build using Buccaneer (Cowtan 2006) followed by manual rebuilding using Coot (Emsley et al., 2010) and further refinement using REFMAC5 (Murshudov et al., 2011) and phenix.refine (Liebschner et al., 2019). The data collection and refinement statistics for all datasets are presented in SI Appendix, Table S2. The final model was deposited in the Protein Data Bank under accession numbers 9FMA (PgPepO) and 9EYG (TfPepO).

## Acknowledgments

These studies were funded by National Science Centre (NCN, Kraków, Poland) Grant Nos. UMO-2018/31/N/NZ1/02891 to I.W. and 2019/35/B/NZ1/03118 to M.K. X-ray data were collected at the BESSY II 14.1 beamline at Helmholtz-Zentrum Berlin für Materialien und Energie (Berlin, Germany) and 11.2C beamline at the Elettra Synchrotron (Trieste, Italy). In addition, we thank the MCB Structural Biology Core Facility (supported by the TEAM TECH CORE FACILITY/2017-4/6 grant from the Foundation for Polish Science) for providing instruments and support.

