## Supplementary Information for "Periodontopathogens interfere with the human renin-angiotensin system through surface-attached proteases"

**Materials and methods**

**Reagents**

High-Capacity cDNA Reverse Transcription Kit, SYBR Green real-time PCR master mix, Hoechst 33342 and Dulbecco's Modified Eagle Medium, DMEM were acquired from Thermo Scientific (Waltham, MA, USA). Vectashield was obtained from VWR (Radnor, PA, USA). Universal RNA Purification Kit was obtained from EURx (Gdańsk, Poland). Hard-Shell® 96-Well PCR Plates, low profile, thin wall and PCR Plate Heat Seal were purchased from Bio-Rad (Hercules, CA, USA).

**Recombinant protein expression and purification**

*Escherichia coli* strain BL21 (DE3) (New England Biolabs, Ipswich, MA, USA) transformed with expression plasmid were grown in LB Lennox media at 37°C to OD_600_ _nm_ ranging from 0.75 to 1.0 and cooled for 30 min at 4°C. Recombinant fusion protein expression was induced with 0.1 mM isopropyl-1-β-d-thiogalactopyranoside (IPTG). After 16 h at 20°C, the cells were collected by centrifugation (6000 × *g*, 10 min, 4°C), resuspended in PBS, pH 7.3 supplemented with 0.02% sodium azide (PBSN), and lysed by sonication (a cycle of 30 × 0.5 s pulses at 70% power output per pellet from 1 L culture) using a Branson digital 450 sonifier (Branson Ultrasonics, Danbury, CT, USA). The cell lysates were clarified by centrifugation (50,000 × *g*, 50 min, 4°C), and recombinant proteins were purified by affinity chromatography on glutathione-Sepharose 4 Fast Flow resins with on-column GST-tag removal by PreScission protease according to manufacturer’s protocol. The final purification step of tag-free recombinant proteins was accomplished by size exclusion chromatography (SEC) on a HiLoad 16/600 Superdex 200 pg pre-equilibrated with 20 mM Tris-HCl, 50 mM NaCl, 0.02% NaN_3_ (pH 8.0). Protein purity during purification was analyzed by SDS-PAGE (Schägger & Jagow, 1987), and protein concentration was taken as the average of two methods: BCA Protein Assay Kit (Thermo Fisher Scientific) and absorbance measurement at 280 nm with a NanoDrop One (Thermo Fisher Scientific) using theoretical extinction coefficient obtained with ProtParam (https://web.expasy.org/protparam/).

**Liquid Chromatography with Tandem Mass Spectrometry (LC-MS/MS)**

Sample desalting was achieved using homemade reverse-phase micro-columns packed with C18 material (3M, [Maplewood](https://www.bing.com/ck/a?!&&p=56540ca15f4ae2acJmltdHM9MTcxODc1NTIwMCZpZ3VpZD0zZWFjYWYzNy01ZGU5LTYxZjItMTgxMC1iYzZiNWM5ZTYwYzEmaW5zaWQ9NTgyNg&ptn=3&ver=2&hsh=3&fclid=3eacaf37-5de9-61f2-1810-bc6b5c9e60c1&u=a1L3NlYXJjaD9GT1JNPVNOQVBTVCZxPU1hcGxld29vZCZmaWx0ZXJzPXNpZDoiYWQ5N2M0MTYtNDBiMS1jYjkyLWI3MDAtMjM2MGVhYjM5MTk3Ig&ntb=1), MN, USA). The LC-MS/MS analysis was conducted on an EASY-nLC 1200 system (Thermo Scientific), interfaced with an Orbitrap Eclipse Tribrid mass spectrometer (Thermo Fisher Scientific), operating in data-dependent acquisition mode. Peptide separation was performed on a 15 cm analytical column with a 75 µm inner diameter, packed in-house with ReproSil-Pur C18-AQ 3 µm resin (Dr. Maisch GmbH, Ammerbuch-Entringen, Germany). The separation utilized a flow rate of 250 nL/min and a 50-minute gradient ranging from 5% to 35% phase B (0.1% formic acid, 90% acetonitrile). The data analysis was carried out using the Mascot search engine, with parameters set for no-enzyme cleavage, cysteine propionamidylation as a fixed modification, and methionine oxidation as a variable modification.

**Analysis of protein binding by PepOs**

Mixtures (final volume: 250 µl) containing the protein: fibrinogen, fibronectin, ferritin or haemoglobin (200 µg) and the PepO at a molar ratio of 1:1 were prepared. Samples were kept at 37°C for 15 min, centrifuged (10 min, 16 900 × g, 4°C) and supernatants were analyzed by SEC on a Superdex 200 increase 10/300 GL column (Cytiva) in 20 mM Tris, 50 mM NaCl, 2.5 mM CaCl_2_, 0.02% NaN_3_ (pH 7.5).

**Confocal microscopy**

Isolated *T. forsythia* cells (C fraction) were divided into two equal parts: one was centrifuged (5 min, 4 500 × g, 4°C), fixed in 1 ml of 3.8% PFA (10 min, 450 rpm, 20°C), while the second one was centrifuged (5 min, 4 500 × g, 4°C), suspended in 1 ml of PBS and left for staining (live cells). Fixed and live cells were washed once with PBS, and divided into 3 parts. To one of them, after centrifugation (5 min, 4 500 × g, 4°C) and removal of the supernatant, 100 µl of polyclonal rabbit anti-TfPepO primary antibodies (at a dilution of 1:50 in PBS) were added and shaken (45 min, 450 rpm, 37°C). Then, the cells were washed once in PBS and suspended in 100 µl of a solution of goat anti-rabbit IgG Alexa Fluor™ 488 diluted 100× in PBS and shaken (45 min, 450 rpm, 37 °C). The cells were washed with PBS, incubated with 100 μl of 0.3 mg/ml Hoechst 33342 solution, centrifuged (10 min, 450 rpm, 20°C) and washed again with PBS (twice). In both cases, cells to which no antibodies were added or to which only a secondary antibody was added were used as negative controls. Drops (20 µl) of bacterial suspensions were placed on microscope slides, "vectashield" was added and analyzed using an LSM880 confocal microscope (100× objective, immersion) and ZEN software (Carl Zeiss, New York, USA).

**Determination of expression of Mas receptor in human gingival fibroblasts**

Total RNA was isolated and reversely transcribed employing the Universal RNA Purification Kit (EURx, Gdańsk, Poland) and the High-Capacity cDNA Reverse Transcription Kit (Thermo Scientific), respectively. PCR reactions were performed at a 20 µL scale with a SYBR Green real-time PCR master mix (Thermo Scientific). Briefly, 4 µl of diluted 200× cDNA was mixed with 5 µl of SYBR Green master mix and primers (0.5 µl each, final concentration: 0.25 µM). The qPCR was performed using CFX Opus 96 Real-Time PCR system (Bio-Rad). Relative quantification was performed in duplicate and by normalizing target gene expression on *EF-2* as housekeeping gene.

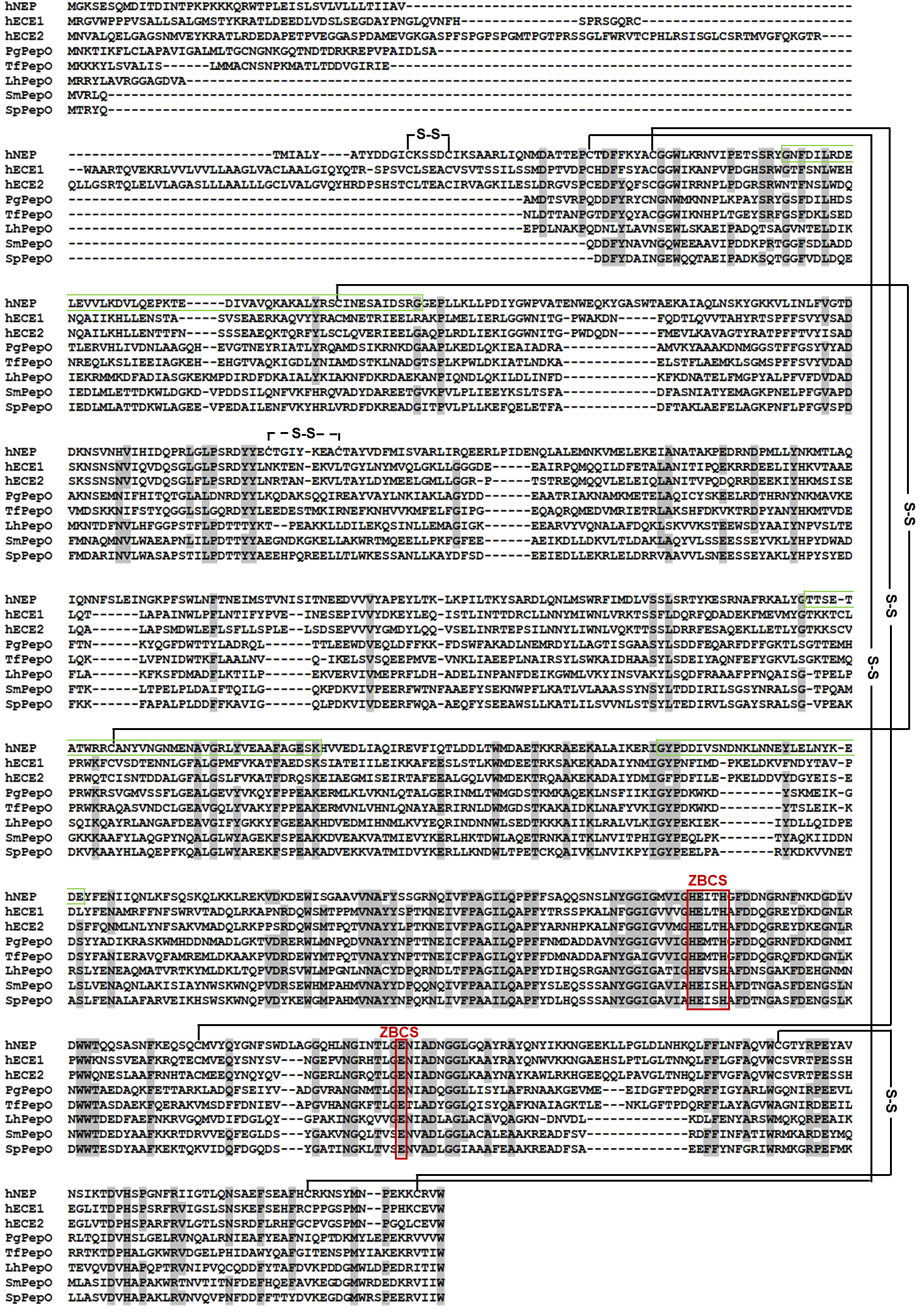

**Fig. S1. Sequence alignment of PgPepO (KEGG database accession number: K07386) and TfPepO (GenBank accession number: KKY61269.1) and other members of the M13 family of proteases: human neprilysin, hNeP, (UniProt accession number, UP no.: P08473); human endothelin convertase-1, hECE1, (UP no.: P42892); human endothelin convertase-2, hECE2, (UP no.: P0DPD6); PepOs from other bacteria: *Lactobacillus helveticus*, LhPepO, (UP no.: O52071); *Streptococcus mutans*, SmPepO, (UP no.: Q8KSS9); and *Streptococcus pneumoniae*, SpPepO (UP no.: A0A4J0IYA2).**

Amino acid residues identical in at least six of the eight aligned sequences are indicated by black letters on a grey background. Zinc-binding consensus sequences (ZBCS) are marked by a red box, while the linker region connecting the regulatory domain and the catalytic domain in neprilysin is indicated with a green frame. Disulfide bridges in hNEP, hECE1, and hECE2 are indicated by black lines (the disulfide bond found only in hNEP is indicated by a dashed black line). Sequence alignment was performed with the Clustal Omega program (https://www.ebi.ac.uk/Tools/msa/clustalo/).

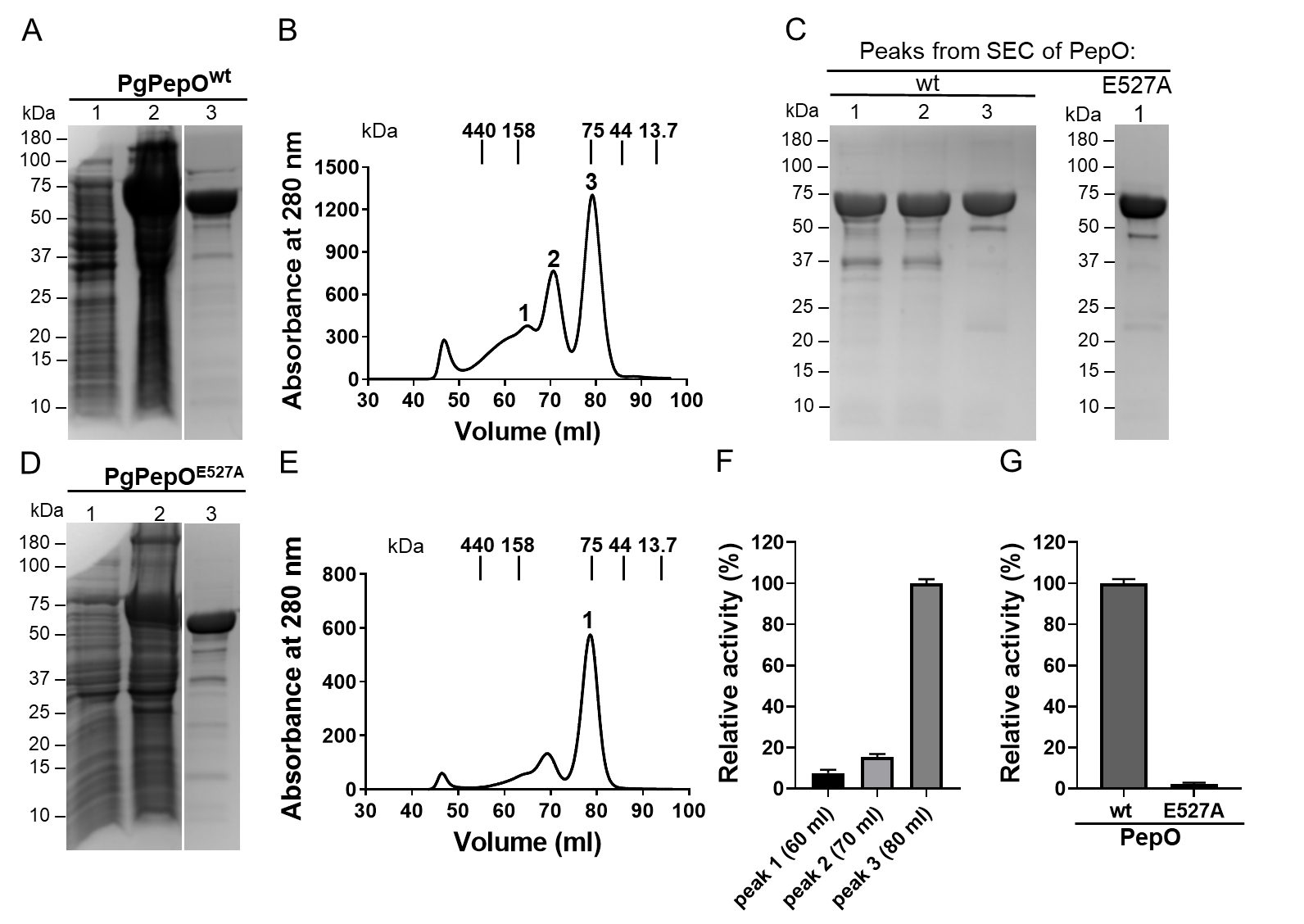

**Fig. S2. Expression and purification of recombinant PgPepO: wild-type (wt) and its catalytic inactive mutant, E527A.**

(A, D) SDS-PAGE analysis of samples obtained during the purification of PgPepO: wt (A) and E527A (D): lanes 1 and 2, *Escherichia coli* cells before and 16 h after induction of protein expression with IPTG, respectively; lane 3, the tag-free PgPepO: wt and E527A (~75 kDa) proteins after purification by affinity chromatography on glutathione-sepharose resins with on-column removal of the GST by cleavage with PreScission protease. (B, E) Chromatograms obtained during further purification of PgPepO: wt (B) and E527A (E) by size exclusion chromatography. The fractions from indicated peaks were pooled and then analyzed by SDS-PAGE (C) and, only for PgPepO^wt^, measurement of activity against Mca-RPPGFSAFK(Dnp)-OH substrate (F). (G) Comparison of PgPepO: wt and E527A activity against Mca-RPPGFSAFK(Dnp)-OH. The activity of peak 3 (F) and PgPepO^wt^ (G) was arbitrarily taken as 100%. The presented data (F, G) are mean±SD (N=3).

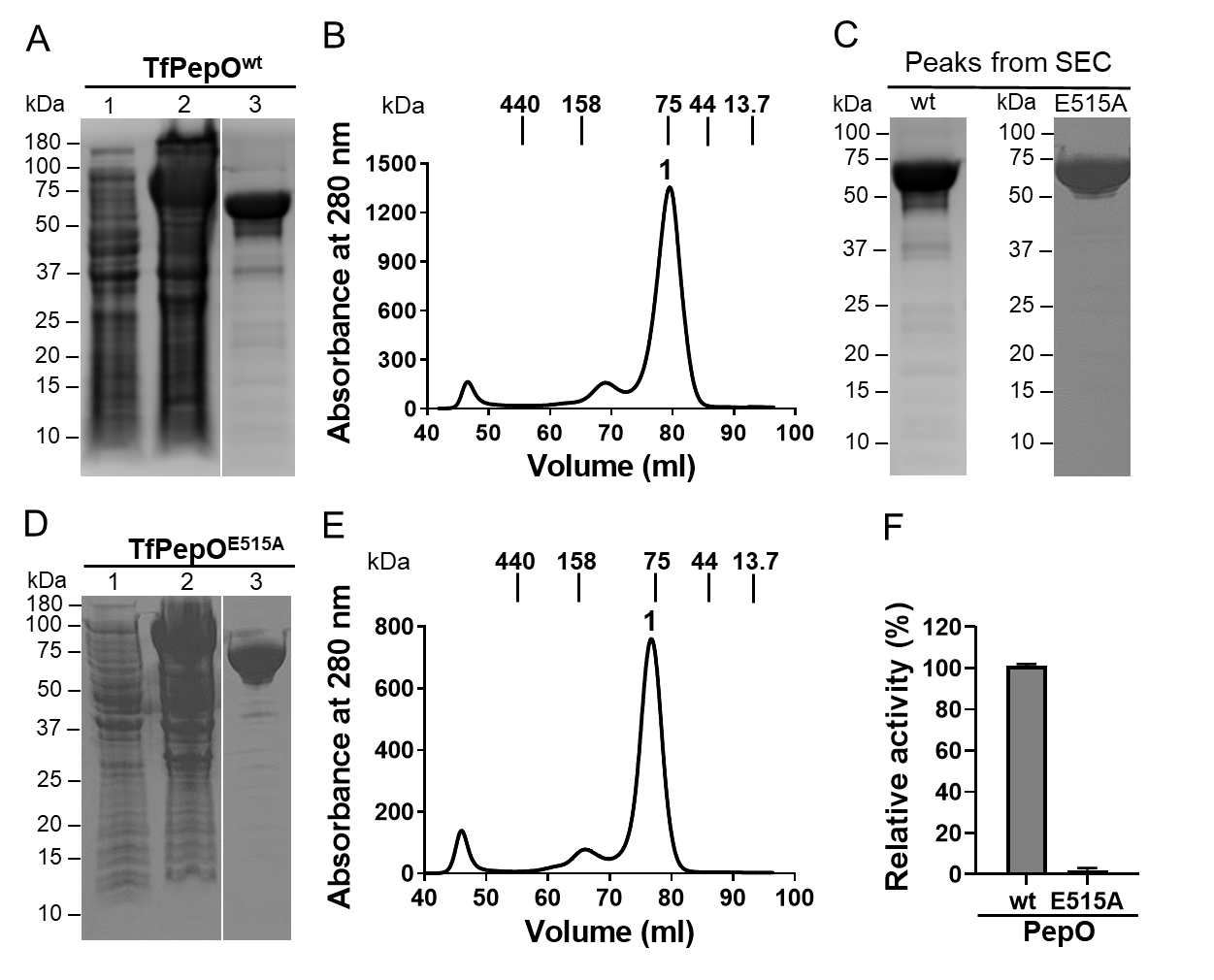

**Fig. S3. Expression and purification of recombinant TfPepO: wild-type (wt) and its catalytic inactive mutant, E515A.**

(A, D) SDS-PAGE analysis of samples obtained during the purification of TfPepO: wt (A) and E515A (D): lanes 1 and 2, *Escherichia coli* cells before and 16 h after induction of protein expression with IPTG, respectively; lane 3, the tag-free TfPepO: wt and E527A (~75 kDa) proteins after purification by affinity chromatography on glutathione-sepharose resins with on-column removal of the GST by cleavage with PreScission protease. (B, E) Chromatograms obtained during further purification of TfPepO: wt (B) and E515A (E) by size exclusion chromatography. The fractions from indicated peaks were pooled and then analyzed by SDS-PAGE (C). (F) Comparison of TfPepO: wt and E515A activity against Mca-RPPGFSAFK(Dnp)-OH. The activity of TfPepO^wt^ was set to 100%. The presented data are mean±SD (N=3).

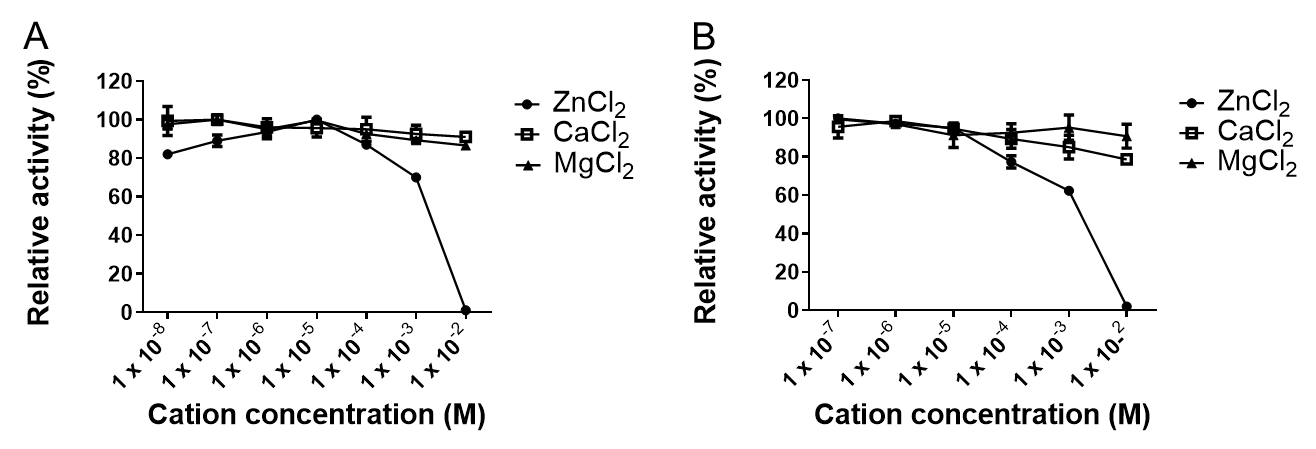

**Fig. S4. Effect of divalent cations on the activity of PgPepO (A) and TfPepO (B).**

PepOs were incubated with increasing concentrations of three cations: Zn^2+^, Ca^2+^ and Mg^2+^ for 15 min in 100 mM Tris, 50 mM NaCl, 0.02% NaN_3_, 0.05% Pluronic F-127 (pH 7.5) and activity against Mca-RPPGFSAFK(Dnp)-OH was measured. Concentrations with the highest activity were arbitrarily taken as 100%. Data shown are mean±SD (N=3).

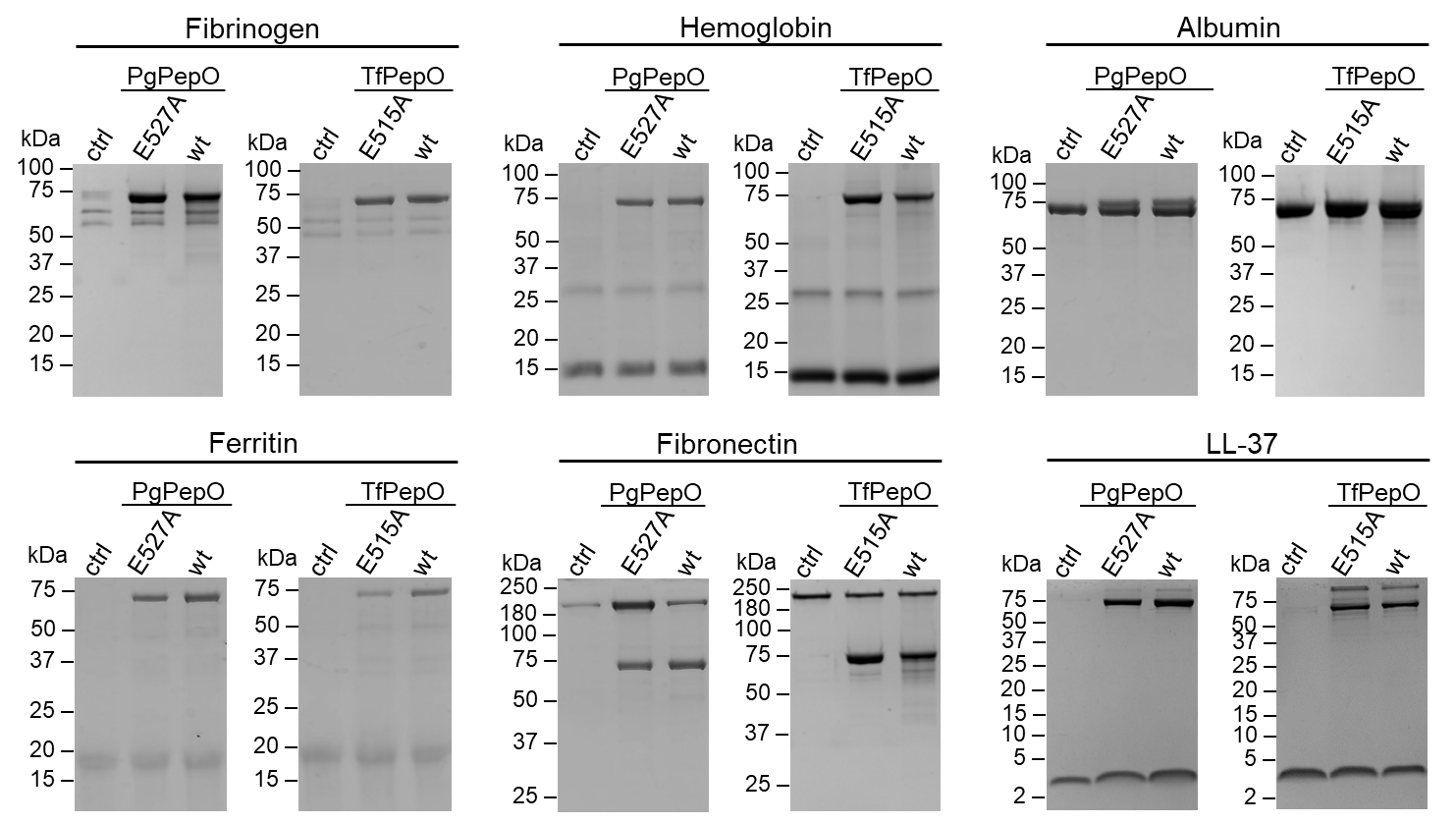

**Fig. S5. Hydrolysis of the peptide and proteinaceous substrates by PgPepO and TfPepO.** Proteases were incubated with substrates at 80:1 substrate:enzyme weight ratio for seven days at 37°C. Obtained samples were analysed by SDS-PAGE. Substrates incubated alone (ctrl) or with appropriate catalytically inactive mutants were used as negative controls.

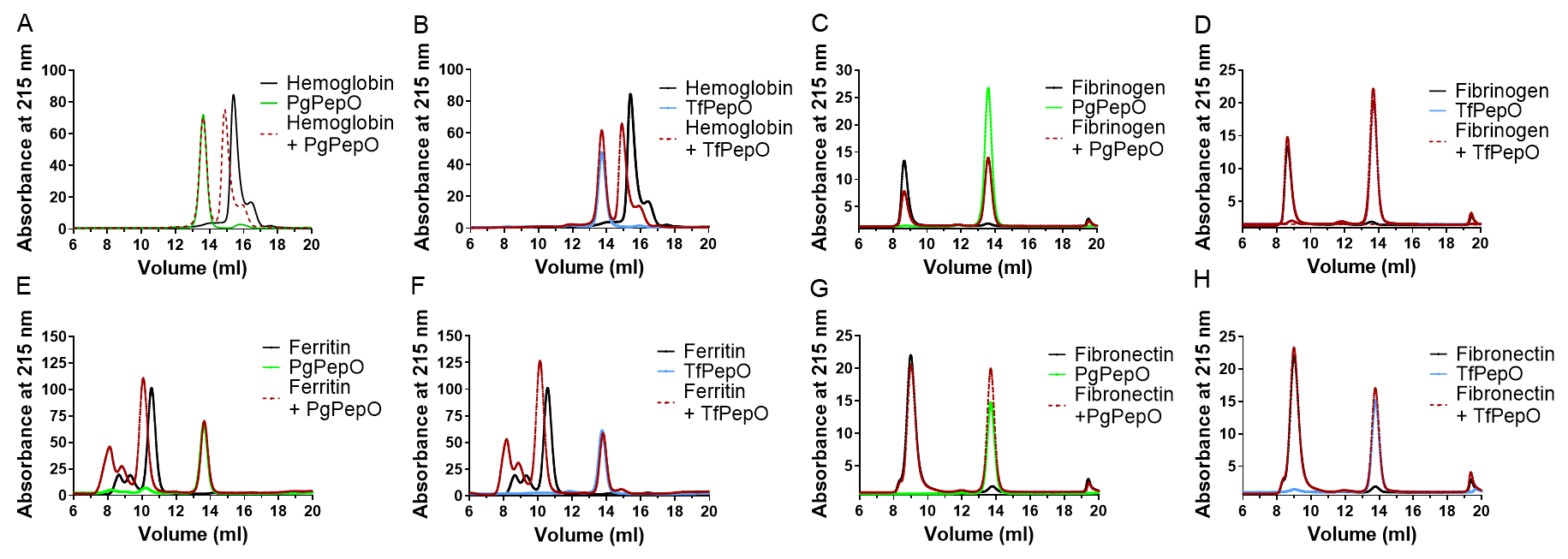

**Fig. S6. Analysis of the binding of various proteins by PgPepO and TfPepO proteases using size exclusion chromatography.**

Chromatograms obtained from the separations of the following protein-protease pairs: hemoglobin-PgPepO (A), hemoglobin-TfPepO (B), fibrinogen-PgPepO (C), fibrinogen-TfPepO (D), ferritin-PgPepO (E), ferritin-TfPepO (F), fibronectin-PgPepO (G) and fibronectin-TfPepO (H). Proteases and proteins were also separated alone.

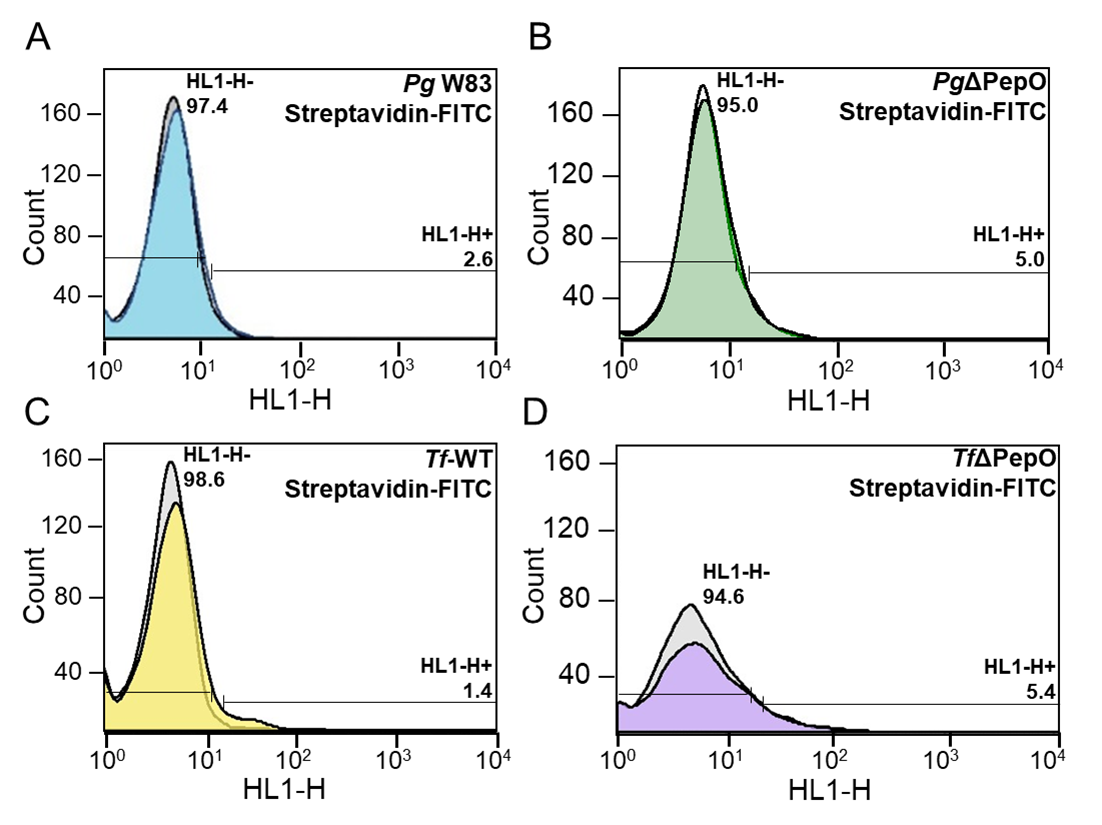

**Fig. S7. Flow cytometry analysis of *P. gingivalis*: W83 (wild-type, wt) and ΔPepO (B), and *T. forsythia*: wt (WT-*Tf*) (C) and ΔPepO (D) to detect biotinylated inner membrane proteins using Streptavidin-Fluorescein Isothiocyanate (FITC) Conjugate.**

Unstained cells are marked with grey histograms.

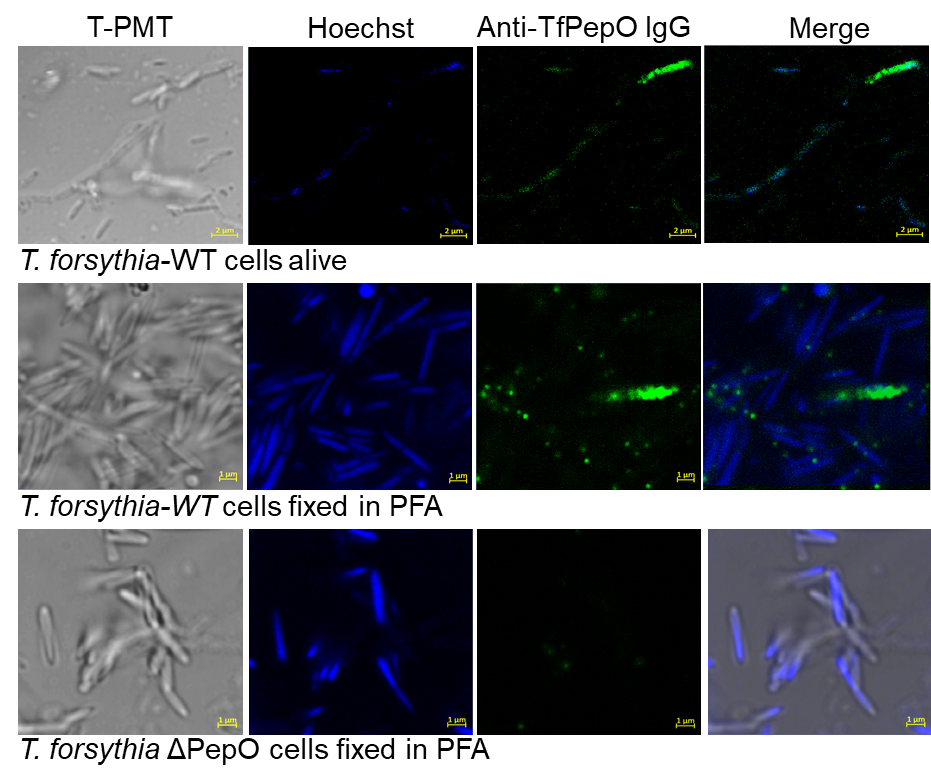

**Fig. S8. Confocal microscopy imaging of TfPepO.**

*T. forsythia* cells: wild-type alive (WT-*Tf*) (upper row), fixed in 3.8% PFA (middle row) and *Tf*ΔPepO (bottom row) were stained using rabbit anti-TfPepO antibody and Hoechst 33342 (detection of genomic DNA) and analyzed under immersion at 100× magnification.

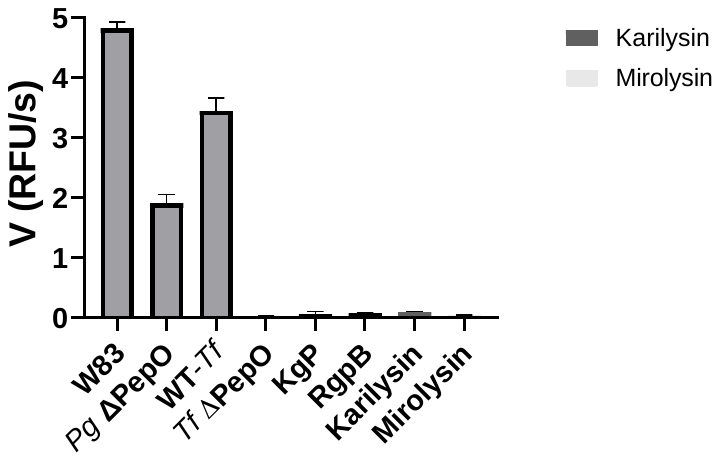

**Fig. S9: Activitity of whole cultures (WC) and purified *P. gingivalis* and *T. forsythia* proteases against the Mca-RPPGFSAFK(Dnp)-OH substrate.**

50 μl of mixture in assay buffer (100 mM Tris, 150 mM NaCl, 2.5 mM CaCl_2_, 0.02% NaN_3_, 0.05% Pluronic F-127 (pH 7.5) containing WC of *P. gingivalis* W83 (wild-type, wt) and *T. forsythia* 43037 (WT-*Tf*) (25 μl of culture normalized to OD_600 nm_ = 0.5) or purified proteases of these bacteria: gingipains Kgp and RgpB (*P. gingivalis*) and KLIKK proteases: karilysin and mirolysin (*T. forsythia*) at concentration 10 nM were mixed with equal volume of substrate (25 μl) and increase in fluorescence (emission/excitation: 320/410 nm) was recorded over time. Values shown are the mean of 3 replicates ± SD.

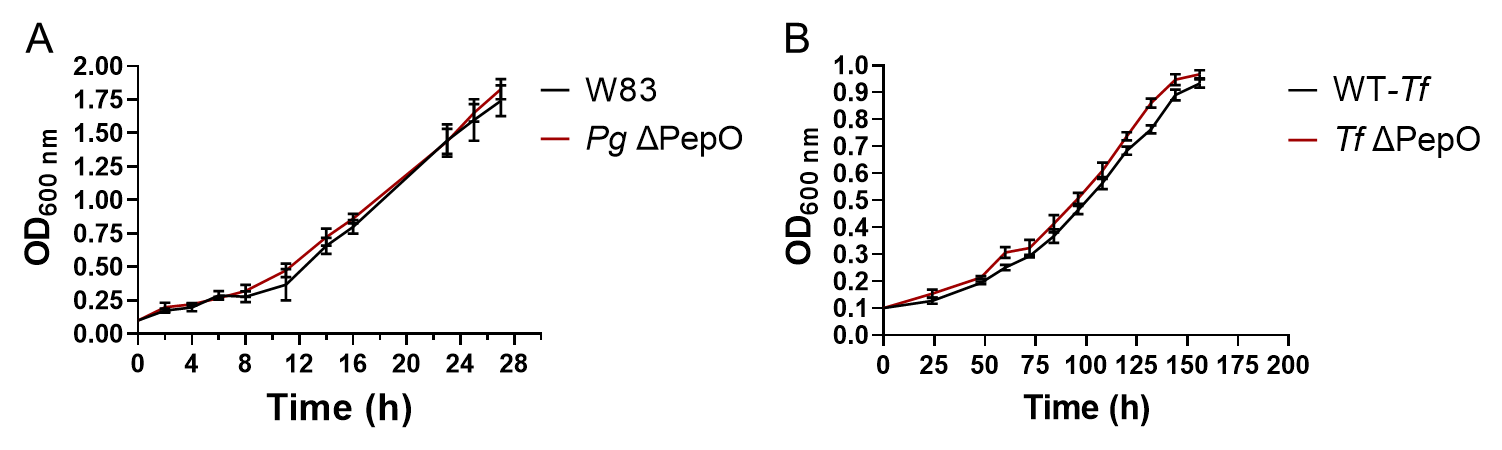

**Fig. S10. Analysis of the effect of PepO deletion on the growth rate of *P. gingivalis* (A) and *T. forsythia* (B).**

*P. gingivalis*: W83 (wild-type, wt) and ΔPepO and *T. forsythia*: WT and ΔPepO bacteria were cultured in minimal medium: DMEM with 2% BSA and OD_600 nm_ values were measured at different time points. Data shown are the mean of 3 replicates ± SD.

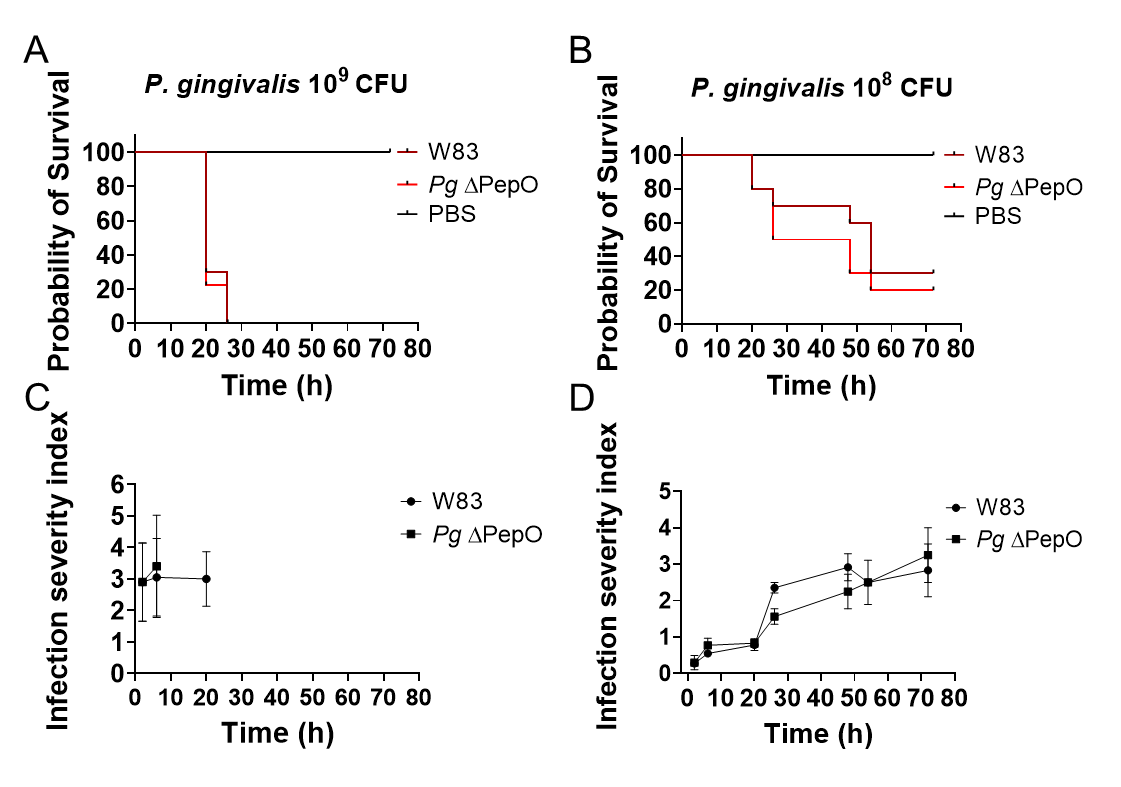

**Fig. S11. Infection of *G. mellonella* larvae by *P. gingivalis*.**

Larvae (N=10 for each group) were infected with two different doses of *P. gingivalis*: W83 (wild-type) and ΔPepO. Dead and live larvae were counted at indicated time points, and the infection severity index was calculated based on the assessment of mobility and melanization of larvae. The data are means±SD and are representative of three biological replicates. Statistical differences were determined by the Kaplan–Meier test (A, B) and one-way ANOVA (C, D).

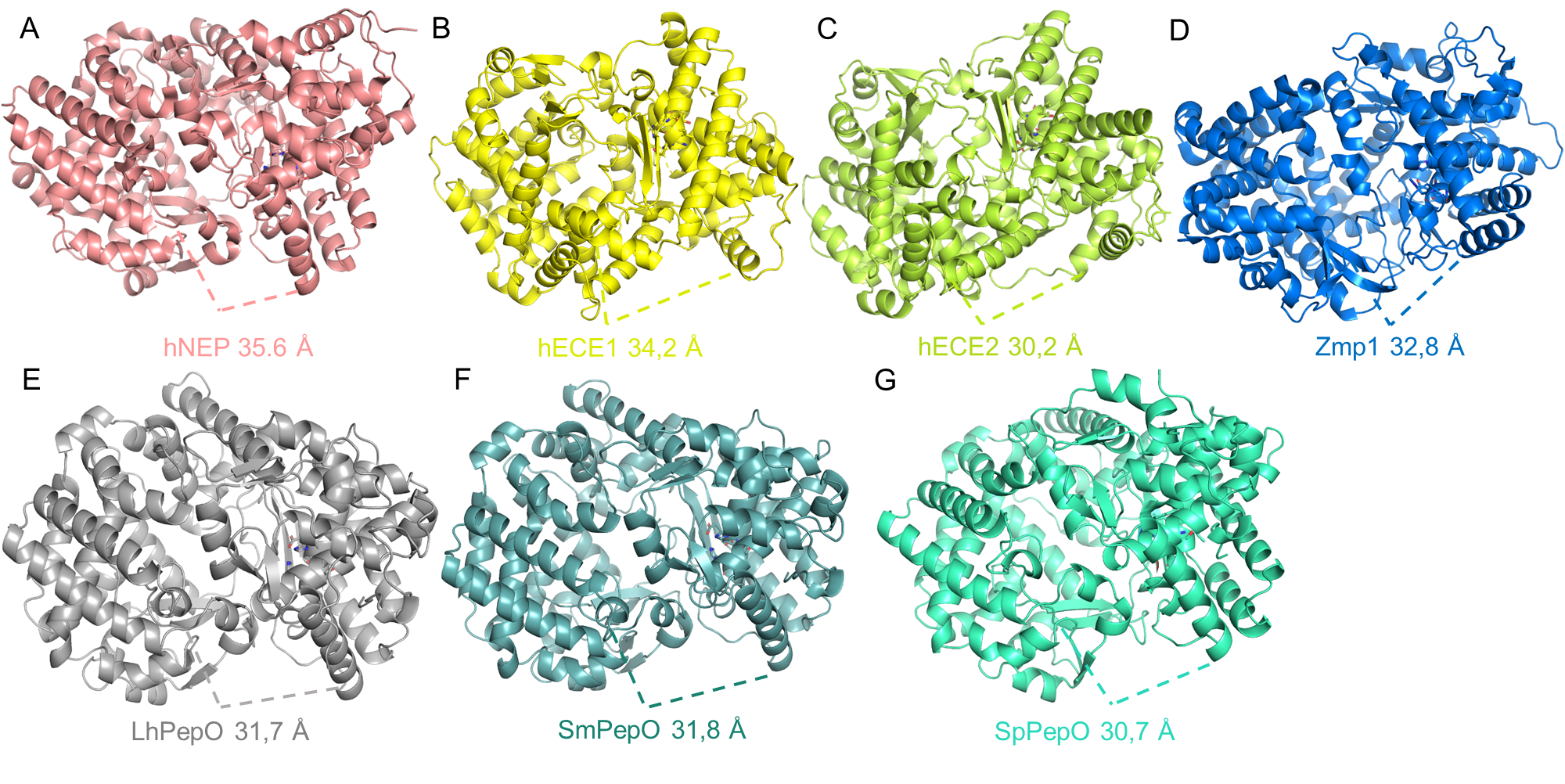

**Fig. S12. General structure of human neprilysin, hNEP (PDB accession number: 6SH1 (A), human endothelin convertase-1, hECE1 (PDB: 3DWB) (B), hECE2 (AlphaFold accession number: AF-P0DPD6-F1), (C), *Lactobacillus helveticus*, LhPepO (PDB: 4IUW) (D), *Streptococcus mutans*, SmPepO (AlphaFold: AF-I6L8Y4-F1), (E), *Streptococcus pneumoniae*, SpPepO (AF-A0A654U8Y8-F1) (F) and *Mycobacterium tuberculosis* PepO, ZMp1 (PDB: 3ZUK) (G).**

Width of the entrance to the catalytic site cleft was measured using PyMol.

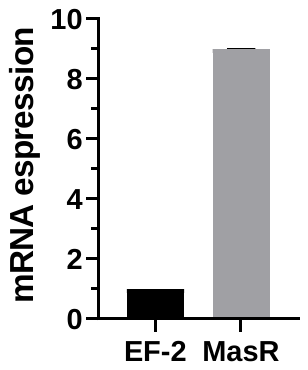

**Fig. S13. Expression of Mas receptor (MasR) in human gingival fibroblasts normalized to the expression of EF-2 as the housekeeping gene.**

The presented data are mean±SD (N=3).

**Table S1. List of primers (5´→3´), strains and plasmids used in this study.**

| **Primers for cloning** | | |
| --- | --- | --- |
| pGEX-6P-1-PgPepO | | |
| PgPepO_F | ACGCGGATCCGCCAATGGCAATAAGGGTCAGACTG | |
| PgPepO_R | ACCGGAATTCTTACCAAACGACTACGCGCTT | |
| **Primers for mutagenesis** | | |
| Mutation | | |
| PgPepO^E527A^_F: | GGCGTAGTGATCGGACACGCGATGACCCATGGATTCGACG | |
| PgPepO^E527A^_R: | CGTCGAATCCATGGGTCATCGCGTGTCCGATCACTACGCC | |
| TfPepO^E515A^_F: | GCGAAAGTGTTTGTGCATGCACTGGGCCATAGC | |
| TfPepO^E515A^_R: | GATCATCAAAGCCATGGGTCATTGCATGGCCAATCACC | |
| **Primers for amplification of genomic fragments containing introduced mutations** | | |
| PgPepOPCR_F | CCTGAATGCTCATGGGCAGGAGCCG | |
| PgPepOPCR_R | CCGAGCCATTGGCAGAAGCCTTTGCG | |
| TfPepOPCR_F | CCTTCCAGTCTATGGTTTCCACATGCC | |
| TfPepOPCR_R | CCAGATTCGACGTGGATGATAAGGCTTC | |
| **Primers for sequencing** | | |
| pGEXF | CCGGGAGCTGCATGTGTCAGAGG | |
| pGEXR | GGGCTGGCAAGCCACGTTTGGTG | |
| PgPepOdsek1_F | GAAACCGATGAAACCCACAC | |
| TfPepOdsek1_F | CACATCTAACTCCGGGACATTG | |
| Erm_R | GATGTTGCAAATACCGATGAGC | |
| Erm_F | CAGGCAAGGGGTTTCTTACTG | |
| PgPepOdsek2_R | CAAGATCGGTGTATTGGTGG | |
| TfPepOdsek2_R | GTCTTACCGCGAATATTGCAC | |
| **Primers for qPCR** | | |
| MasR_F | AGGGTGACTGACTGAGTTTGG | |
| MasR_R | GAAGGTAAGAGGACAGGAGC | |
| EF-2_F | GACATCACCAAGGGTGTGCAG | |
| EF-2_R | TTCAGCACACTGGCATAGAGGC | |
| **Bacterial strains** | | |
| **Name** | **Description (genotype, resistance)** | **Source / reference** |
| *Escherichia coli* DH5α | Genotype: F- 80dlacZ M15 (lacZYA-argF) U169 recA1 endA1hsdR17(rk-, mk+) phoAsupE44 -thi-1 gyrA96 relA1 | Thermo Fisher Scientific |
| *Escherichia coli* BL21 (DE3) | A strain lacking two peptidases: lon and ompT | Merck-Millipore |
| *Porphyromonas gingivalis* W83 | Wild-type strain (WT) | Department Microbiology Collection |
| *Porphyromonas gingivalis* ΔPepO (*Pg* ΔPepO) | Wild-type strain W83 with deletion of the PgPepO gene, resistant to erythromycin | This study |
| *Tannerella forsythia* ATCC 43037 | Wild-type strain (WT) | ATCC 43037 |
| *Tannerella forsythia* ΔPepO (*Tf* ΔPepO) | Wild-type strain ATCC 43037 with deletion of the TfPepO gene, resistant to erythromycin | This study |
| **Plasmids** | | |
| pGEX-6P-1-PgPepO | Expression construct of the PgPepO protein with an N-terminal GST tag | This study |
| pGEX-6P-1-TfPepO | Expression construct of the TfPepO protein with an N-terminal GST tag | This study |
| pGEX-6P-1-PgPepO^E527A^ | Expression construct of the PgPepO^E527A^ protein with an N-terminal GST tag | This study |
| pGEX-6P-1-TfPepO^E515A^ | Expression construct of the TfPepO^E515A^ protein with an N-terminal GST tag | This study |
| pUC19-ΔPgPepO | Plasmid for PgPepO deletion in the *P. gingivalis* genome | This study |
| pUC19-ΔTfPepO | Plasmid for TfPepO deletion in the *T. forsythia* genome | This study |
| **Erythromycin resistance cassette sequence** | | |
| ATAGCTTCCGCTATTGCTTTTTTGCTCATCGGTATTTGCAACATCATAGAAATTGCATACCTTTGTTCCTCGGTTATATGTTTGCTCATCTGCAACTTTTTTTTCTTTGGACGGACAATTAAAGCAAAGATAGCAAACTTTATCCATTCAGAGTGAGAGAAAGGGGGACATTGTCTCTCTTTCCTCTCTGAAAAATAAATGTTTTTATTGCTTATTATCCGCACCCAAAAAGTTGCATTTATAAGTTGAACTCAAGAAGTATTCACCTGTAAGAAGTTACTAATGACAAAAAAGAAATTGCCCGTTCGTTTTACGGGTCAGCACTTTACTATTGATAAAGTGCTAATAAAAGATGCAATAAGACAAGCAAATATAAGTAATCAGGATACGGTTTTAGATATTGGGGCAGGCAAGGGGTTTCTTACTGTTCATTTATTAAAAATCGCCAACAATGTTGTTGCTATTGAAAACGACACAGCTTTGGTTGAACATTTACGAAAATTATTTTCTGATGCCCGAAATGTTCAAGTTGTCGGTTGTGATTTTAGGAATTTTGCAGTTCCGAAATTTCCTTTCAAAGTGGTGTCAAATATTCCTTATGGCATTACTTCCGATATTTTCAAAATCCTGATGTTTGAGAGTCTTGGAAATTTTCTGGGAGGTTCCATTGTCCTTCAATTAGAACCTACACAAAAGTTATTTTCGAGGAAGCTTTACAATCCATATACCGTTTTCTATCATACTTTTTTTGATTTGAAACTTGTCTATGAGGTAGGTCCTGAAAGTTTCTTGCCACCGCCAACTGTCAAATCAGCCCTGTTAAACATTAAAAGAAAACACTTATTTTTTGATTTTAAGTTTAAAGCCAAATACTTAGCATTTATTTCCTGTCTGTTAGAGAAACCTGATTTATCTGTAAAAACAGCTTTAAAGTCGATTTTCAGGAAAAGTCAGGTCAGGTCAATTTCGGAAAAATTCGGTTTAAACCTTAATGCTCAAATTGTTTGTTTGTCTCCAAGTCAATGGTTAAACTGTTTTTTGGAAATGCTGGAAGTTGTCCCTGAAAAATTTCATCCTTCGTAGTTCAAAGTCGGGTGGTTGTCAAGATGATTTTTTTGGTTTGGTGTCGTCTTTTTTTAAGCTGCCGCATAACGGCTGGCAAATTGGCGATGGAGCGGAAAC | | |

**Table S2. Statistics for X-ray crystallographic data collection and refinement.**

|  | **PgPepO** | **TfPepO** |
| --- | --- | --- |
| PDB code | 9FMA | 9EYG |
| Synchrotron / beamline | HZB BESSY / 14.1 | Elettra Synchrotron / 11.2C |
| Data collection | | |
| Space group | P 41 21 2 | P 3 2 2 1 |
| Cell dimensions  a, b, c (Å)  α, β, γ (º) | 93.2 93.2 282.5  90 90 90 | 95.4 95.4 301.9  90 90 120 |
| Unique reflections | 80700 (5260) | 148384 (4837) |
| Resolution range (Å) | 45.16-1.76 (1.86-1.76) | 48.7 -1.8 (1.86-1.80) |
| Completeness (%) | 99.9 (99.6) | 99.96 (99.99) |
| CC(1/2) | 99.9 (38.7) | 99.6 (60.5) |
| I/σI | 13.02 (0.93) | 16.92 (1.52) |
| Redundancy | 14.4 | 9.3 |
| Wilson B-factor | 27.81 | 29.81 |
| Refinement |  |  |
| Resolution range (Å) | 46.60-1.75 | 48.70-1.80 |
| Rwork / Rfree | 0.18 / 0.21 | 0.19 / 0.22 |
| Number of non-hydrogen atoms Protein  Solvent  Ligands | 5289  506  5 | 10583  804  153 |
| R. m. s deviations  bonds (Å) angles (°) | 0.005  0.66 | 0.009  0.97 |
| Average B-factor (Å^2^) | 31.92 | 35.59 |
| Ramachandran plot (%)  Favored  Allowed  Outliers | 98.31  1.69  0.00 | 98.38  1.62  0.00 |

Values in parentheses are for highest-resolution shell.
